# Genome organization regulates nuclear pore complex formation and promotes differentiation during Drosophila oogenesis

**DOI:** 10.1101/2023.11.15.567233

**Authors:** Noor M. Kotb, Gulay Ulukaya, Ankita Chavan, Son C. Nguyen, Lydia Proskauer, Eric Joyce, Dan Hasson, Madhav Jagannathan, Prashanth Rangan

## Abstract

Genome organization can regulate gene expression and promote cell fate transitions. The differentiation of germline stem cells (GSCs) to oocytes in Drosophila involves changes in genome organization mediated by heterochromatin and the nuclear pore complex (NPC). Heterochromatin represses germ-cell genes during differentiation and NPCs anchor these silenced genes to the nuclear periphery, maintaining silencing to allow for oocyte development. Surprisingly, we find that genome organization also contributes to NPC formation, mediated by the transcription factor Stonewall (Stwl). As GSCs differentiate, Stwl accumulates at boundaries between silenced and active gene compartments. Stwl at these boundaries plays a pivotal role in transitioning germ-cell genes into a silenced state and activating a group of oocyte genes and Nucleoporins (Nups). The upregulation of these Nups during differentiation is crucial for NPC formation and further genome organization. Thus, crosstalk between genome architecture and NPCs is essential for successful cell fate transitions.

## Introduction

During oogenesis, germline stem cells (GSCs) differentiate and undergo meiosis to generate oocytes.^1–4^ These oocytes accrue a maternally synthesized trust fund of RNAs called maternal RNAs required to launch the next generation.^4–7^ Drosophila oogenesis has a well-characterized transition from GSC to oocyte (**Figure 1A-A1**).^3,6,8–14^ A programmatic transition, referred to as the germ cell-to-maternal transition (GMT), promotes the silencing of genes that are expressed during the early stages of oogenesis, including a cohort of differentiation-promoting genes (**Figure 1A1)**.^15,16^ These “early oogenesis genes’’ include *ribosomal small subunit protein19b* (*rpS19b*) and *blanks*.^15–17^ The transcriptional silencing of these early oogenesis genes is mediated by SET Domain Bifurcated Histone Lysine Methyltransferase 1 (SETDB1), which leads to methylation of H3K9 and thereby establishes heterochromatin.^15,18,19^ Once transcriptionally silenced, these heterochromatic regions containing early oogenesis genes are anchored to the nuclear periphery mediated by the nuclear pore complex (NPC).^15,20–22^ Loss of SETDB1 or specific nucleoporins (Nups) that comprise the NPCs leads to the upregulation of early oogenesis genes in the egg chambers that fail to grow and result in sterility.^15,18,23^ Silencing of these early oogenesis genes is concomitant with increased expression of maternally deposited genes.^15,16^ The mechanism by which these large gene expression changes are regulated during this transition is not fully understood.

**Figure1.**
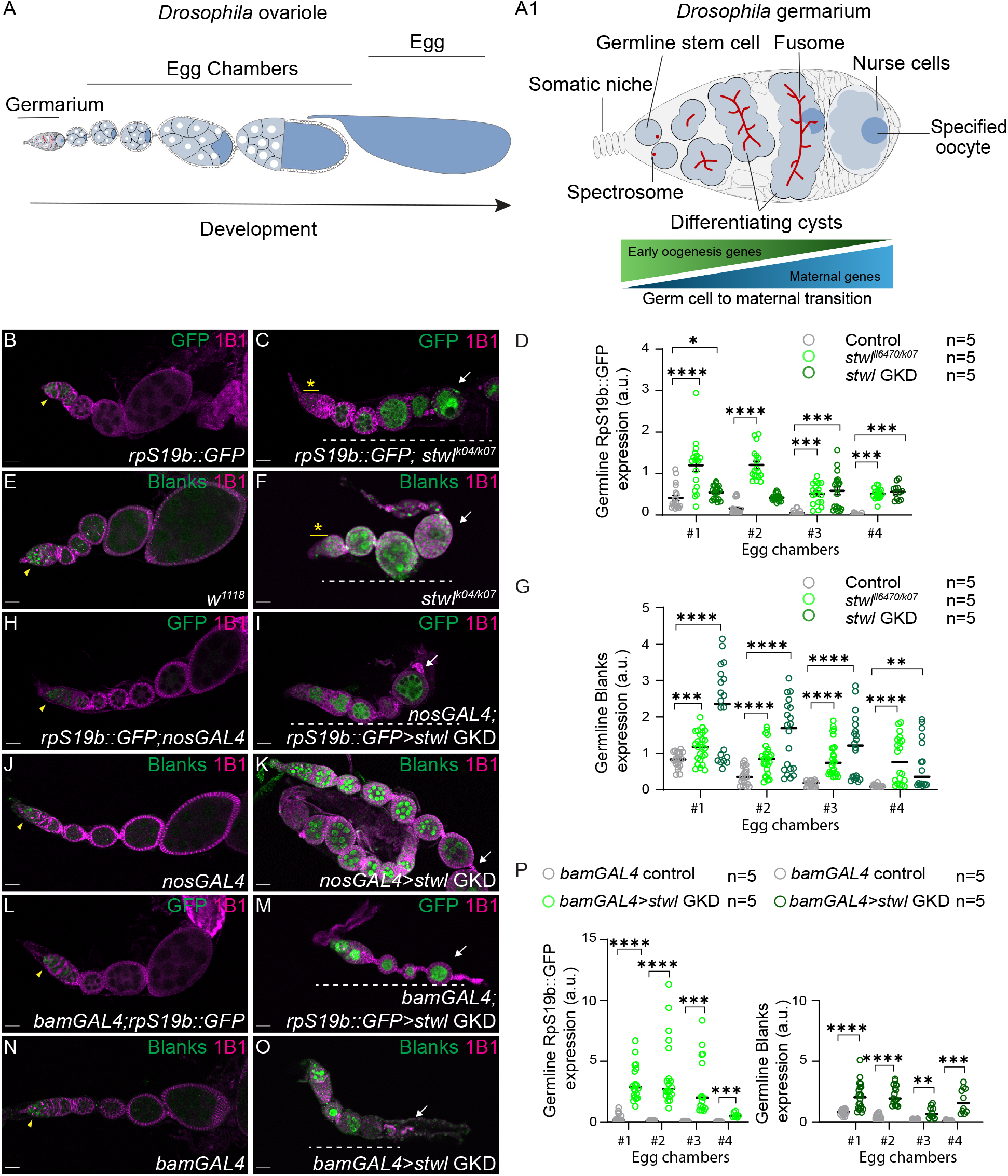
*stwl* is required for silencing RpS19b::GFP and Blanks during oogenesis. (A) A schematic of a *Drosophila* ovariole consisting of a germarium and egg chambers surrounded by somatic cells (white). Egg chambers grow and produce an egg (dark blue). (A1) A schematic of a *Drosophila* germarium. Germline stem cells (GSCs; blue) are proximal to somatic niche (grey) and divide to give rise to daughter cells called cystoblasts. Both GSCs and cystoblasts are marked by spectrosomes (red). Cystoblasts differentiate, giving rise to 2-, 4-, 8-, and 16-cell cysts (blue), marked by fusomes (red). In the 16-cell cyst, one cell commits to meiosis and specifies an oocyte (dark blue), whereas the other 15 cells become nurse cells (light blue). Early oogenesis genes are expressed in the undifferentiated stages and are attenuated upon oocyte specification while maternal genes increase and enriched in the differentiated stages. This transition happens in a programmatic manner called the germ cell to maternal transition (GMT). (B–C) Ovariole of control (B) and *stwl^K^*^04^*^/K^*^07^ carrying *rpS19b::GFP* (C) stained for GFP (green) and 1B1 (magenta). In control, GFP is expressed in the undifferentiated stages and early cysts and then silenced (yellow arrows). In *stwl* mutants, egg chambers ectopically expressed RpS19b::GFP (white dashed line), did not grow, and died during oogenesis. Yellow asterisk indicates loss of germline stem cells (GSCs) in *stwl^K^*^04^*^/K^*^07^. (D) Arbitrary unit (a.u.) quantification of RpS19b::GFP expression in egg chambers in *stwl^k^*^04^*^/k^*^07^ and upon the GKD of *stwl* (green) compared to control ovaries (grey). In the control, GFP is expressed in the undifferentiated cells and is attenuated in the egg chambers. In *stwl^k^*^04^*^/k^*^07^ and *stwl* GKD, GFP expression persists in the egg chambers. Statistics: Two tailed t-test; n = 5 ovarioles per genotype; ns, P > 0.05; ∗P< 0.05; ∗∗P < 0.01; ∗∗∗P < 0.001; ∗∗∗∗P < 0.0001. (E-F) Ovariole of control *w*^1118^ (E) and *Stwl^K^*^04^*^/K^*^07^ (F) stained for Blanks (green) and 1B1 (magenta). Blanks is expressed in the undifferentiated cells (yellow arrows) and is attenuated in the egg chambers. *stwl* mutants resulted in egg chambers that ectopically expressed Blanks (white dashed line), did not grow, and died during oogenesis. Yellow asterisk indicates loss of germline stem cells (GSCs) in *stwl ^K^*^04^*^/K^*^07^. (G) Arbitrary unit (a.u.) quantification of Blanks expression in egg chambers in *stwl^k^*^04^*^/k^*^07^ and upon the GKD of *Stwl* (green) compared with control ovaries (grey). In the control, Blanks is expressed in the undifferentiated cells and are attenuated in the egg chambers. In *stwl^k^*^04^*^/k^*^07^ and *stwl* GKD, Blanks expression persists in the egg chambers. Statistics: Two tailed t-test; n = 5 ovarioles per genotype; ns, P > 0.05; ∗P < 0.05; ∗∗P < 0.01; ∗∗∗P < 0.001; ∗∗∗∗P < 0.0001. (H–I) Ovariole of control *rpS19b::GFP;nosGAL4* (H) and GKD of *stwl* (I) stained for GFP (green) and 1B1 (magenta).In the control, GFP is expressed in the undifferentiated cells (yellow arrows) and is attenuated in the egg chambers. *stwl* GKD resulted in egg chambers that ectopically expressed RpS19b::GFP (white dashed line), did not grow, and died during oogenesis. (J-K) Ovariole of control (J) and GKD of *stwl* (K) stained for Blanks (green) and 1B1 (magenta). In the control, Blanks is expressed in the undifferentiated cells (yellow arrows) and is attenuated in the egg chambers. *stwl* GKD resulted in egg chambers that ectopically expressed blanks (white dashed line), did not grow, and died during oogenesis. (L-M) Ovariole of control *bamGAL4*;*rpS19b::GFP* (L) and *bamGAL4*;*rpS19b::GFP>stwl* GKD (M) stained for GFP (green) and 1B1 (magenta). In the control, GFP is expressed in the undifferentiated cells (yellow arrows) and is attenuated in the egg chambers. *stwl* GKD in the cyst stages using *bamGAL4*;*rpS19b::GFP* resulted in egg chambers that ectopically expressed RpS19b::GFP (white dashed line), did not grow, and died during oogenesis. (N-O) Ovariole of control (N) and GKD of *stwl* in the cyst stages using *BamGAL4* (O) stained for Blanks (green) and 1B1 (magenta). In the control, Blanks is expressed in the undifferentiated cells (yellow arrows) and is attenuated in the egg chambers. *stwl* GKD in the cyst stages resulted in egg chambers that ectopically expressed blanks (white dashed line), did not grow, and died during oogenesis. (P) Arbitrary unit (a.u.) quantification of early oogenesis proteins RpS19b::GFP and Blanks expression in egg chambers upon the GKD of *stwl* (green) compared with control ovaries (grey). In the control, GFP and Blanks are expressed in the undifferentiated cells and are attenuated in the egg chambers. In *stwl* GKD in the cyst stages, GFP and Blanks expression persist in the egg chambers. Statistics: Two tailed t-test; n = 5 ovarioles per genotype; ns, P > 0.05; ∗P < 0.05; ∗∗P < 0.01; ∗∗∗P < 0.001; ∗∗∗∗P < 0.0001. Scale bars: 15 μm.

Changes in global genome organization play critical roles in regulating differentiation and gene expression.^24–29^ The genome is partitioned into compartments mediated by the formation of conserved topological associated domains (TADs).^30–34^ Each TAD contains genes and regulatory elements that interact more frequently with each other than with regions outside of the TAD.^30,33–35^ TADs are demarcated by boundary elements that interact with insulator proteins, such as cohesin, CTCF, and BEAF, as well as various transcription factors.^36–38^ TADs can regulate gene expression by facilitating enhancer-promoter interactions within their boundaries while insulating genes from neighboring regulatory elements.^24,27,34,39–43^ Lamin-associated domains are TADs that are associated with the nuclear periphery^44,45^ and often contain genes that are transcriptionally silenced or exhibit low expression, which are marked by repressive chromatin marks, including methylated H3K9.^46–48^ Early oogenesis genes such as *rpS19b* are silenced and tethered to the nuclear periphery but how these genes are recruited to the nuclear periphery is not known.

The transcription factor Stonewall (Stwl) is required for Drosophila oogenesis^49–52^, accumulates at insulator elements, and interacts with heterochromatin components such as HP1.^49–51^ Intriguingly, loss of *stwl* phenocopies loss of *SETDB1* or *Nups,* resulting in egg chambers that fail to grow, leading in sterility.^15,18,23,50,51,53^ However, how Stwl regulates oogenesis was not fully known.^49–51^ Here, we discovered that *stwl*, like *SETDB1* and components of the NPC, represses early oogenesis genes during differentiation but is also critical for activating a cohort of maternally supplied genes. We show that Stwl coordinates gene expression changes at the GMT and promotes differentiation by stabilizing chromatin boundaries and promoting tethering of silenced genes to the nuclear periphery. ^42,43,48,54^

## Results

### Stonewall is required during oocyte specification to silence reporter of *rpS19b* and Blanks in the egg chambers

We hypothesized that *stwl*, like *SETDB1*^15^, could regulate the expression of early oogenesis genes. To test this hypothesis, we generated control and *stwl* mutant flies carrying a reporter of *rpS19b, rpS19b::GFP* that is under endogenous control.^16^ *stwl* mutant ovaries lose stem cells and contain egg chambers that do not grow, consistent with previous data (**Figures S1A-S1B)**.^50–52^ We stained ovaries of control and *stwl* mutants carrying the *rpS19b::GFP* for GFP and 1B1, which marks the somatic cell membranes, spectrosomes, and fusomes in the germline.^55^ We also independently stained control and *stwl* mutant ovaries for another early oogenesis protein, Blanks, along with 1B1.^17,56,57^ In contrast to control ovaries where RpS19b::GFP and Blanks are silenced in the differentiated egg chambers, in *stwl* mutants, we found that RpS19b::GFP and Blanks are ectopically expressed in the egg chambers (**Figures 1B-G and Figure S1C)**.^58^ Thus, *stwl* is required for silencing both RpS19b::GFP and Blanks expression in the egg chambers.

Using an antibody raised against Stwl, we found that Stwl is expressed both in the germline and the soma of the ovary **(Figures S1D-S1D1)**. This expression is attenuated in *stwl* mutants **(Figures S1E-S1E1)**. To determine if germline expression of *stwl* is required to promote the silencing of early oogenesis genes, we used a germline-specific driver, *nosGAL4,* to drive RNAi to deplete *stwl* in the germline of *RpS19b::GFP* reporter flies.^59^ Germline knockdown (GKD) of *stwl* resulted in egg chambers that do not grow, phenocopying *stwl* mutants **(Figures S1A-S1B)** and in ectopic RpS19b::GFP and Blanks expression in egg chambers, similar to *stwl* mutants **(Figures 1H-1K and 1G).** Loss of *stwl* in the soma using a gonadal somatic cell-specific driver *tjGAL4* does not result in a phenotype nor does it upregulate Blanks in the egg chambers **(Figures S1F-S1G)**.^60^ From these data, we infer that Stwl is required in the germline to promote the silencing of RpS19b::GFP and Blanks in the egg chambers.

Stwl levels increase in the cyst stages of differentiation **(Figures S1H-S1I)**. Early oogenesis genes, *rpS19b,* and *blanks,* are silenced starting in the cyst stages mediated by SETDB1-dependent heterochromatin formation and then maintained in a silenced state in the egg chambers.^15^ To determine if Stwl is required in the cyst stages during the initiation of silencing or in the later stages for the maintenance of the silenced state, we depleted *stwl* in the cyst stages using *bamGAL4* and in differentiated egg chambers using *Mat*α*GAL4*.^61,62^ Compared to the control, we found that loss of *stwl* in cyst stages resulted in egg chambers that do not grow and express RpS19b::GFP and Blanks **(Figures 1L-1P and Figures S1B)**. In contrast, *stwl* depletion using *Mat*α*GAL4* did not result in a phenotype or upregulation of a RpS19b::GFP (**Figures S1J-S1K**). Taken together, we find that *stwl* is required during differentiation in cyst stages for egg chamber growth and for silencing the RpS19b::GFP and Blanks.

### Stonewall is required to silence early oogenesis genes and activate a cohort of maternal genes

To determine if *stwl* regulates other early oogenesis genes, we conducted RNA-sequencing (RNA-seq) of control and *stwl* GKD ovaries. Importantly, we analyzed both *nosGAL4*-mediated and *bamGAL4-*mediated GKD of *stwl* **(Figure 2A**, **Table 1)**. We used a 2-fold cut-off (Fold Change −2>(Log2FC)>2) and adjusted p-value<0.05 to identify significantly dysregulated genes. The overlapping set of genes in the two GKD ovaries reports on Stwl-regulated genes in the cyst stages of the germline. We use this overlapping set of genes dysregulated in both *nosGAL4-* and *bamGAL4-* mediated *stwl* depletion for all further analysis.

**Figure 2.**
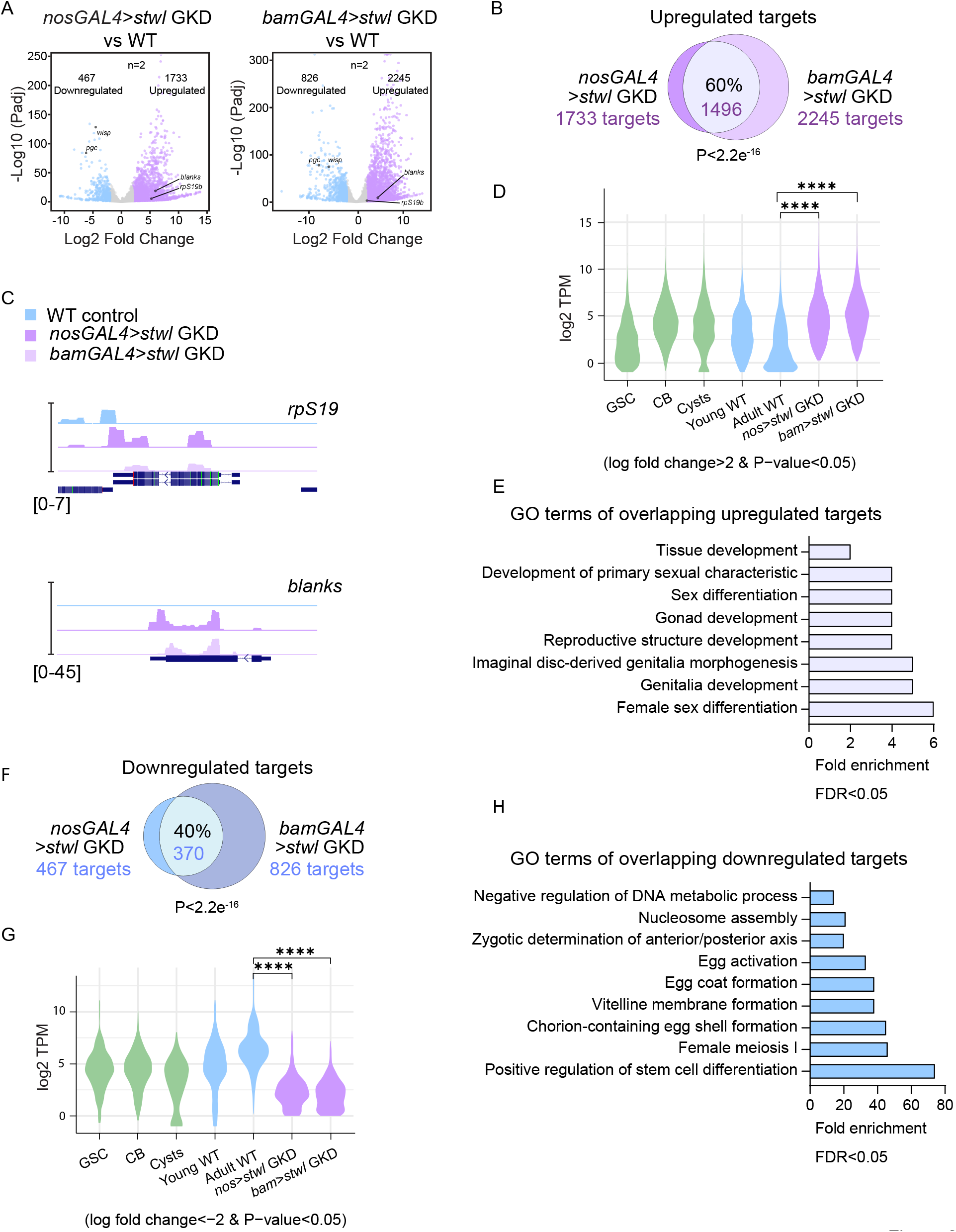
Stwl regulates the silencing of early oogenesis genes and activation of some maternal genes. (A) Volcano plots of −log10p value vs. log2fold change (FC) of *nosGAL4*>*stwl* GKD ovaries vs *nosGAL4* (control) as well as *bamGAL4*>*stwl* GKD ovaries vs *bamGAL4* (cyst control) ovaries showing significantly downregulated (blue) and upregulated (lilac) transcripts in *stwl* GKD ovaries compared to control ovaries (Adjusted p-value< 0.05 and genes with 2-fold or higher change were considered significant). n=2. (B) Venn diagram of upregulated genes from RNA-seq of *nosGAL4 > stwl* GKD ovaries and *bamGAL4> stwl* GKD compared with controls. 60% of targets are shared between both, suggesting that *stwl* is required in the cyst stages to upregulate a cohort of genes. Statistics: hypergeometric test p<2.2e^-16^. (C) RNA-seq tracks showing that *rpS19b* and *blanks* are upregulated upon *stwl* GKD (*nosGAL4)* in dark purple and in the cyst stages (*bamGAL4)* in light purple compared to control (blue). (D) Violin plot of RNA levels of the shared upregulated targets (*nosGAL4*>*stwl* GKD *& bamGAL4>stwl* GKD) in ovaries enriched for GSCs, cytoblasts, cysts, and whole ovaries, showing that Stwl upregulated targets are expressed up to the cyst stages and attenuated in whole ovaries. Statistics: A negative binomial regression model was used to estimate the average TPM count of ‘signature’ genes between each ‘genotype’. The TPM of each gene was used as the dependent variable and ‘genotype’ was the independent variable. Statistical comparisons between groups were performed using contrasts and p-values were adjusted for multiple comparisons using the Benjamini-Hochberg procedure false discovery rate (p-FDR). The average TPM between groups was considered to be significantly different when p-FDR < 0.05. (E) The biological process GO terms of shared upregulated genes in ovaries of *nosGAL4*>*stwl GKD and bamGAL4>stwl* GKD compared with controls using fold enrichment. Statistics: Fisher’s exact test, showing that the overlapping upregulated targets are involved in gonad development and female sex differentiation. Displaying results for FDR< 0.05. (F) Venn diagram of downregulated genes from RNA-seq of *nosGAL4 > stwl* GKD ovaries and *bamGAL4> stwl* GKD compared with controls. 40% of targets are shared between both, suggesting that Stwl is required in the cyst stages to upregulate a cohort of genes. Statistics: hypergeometric test P<2.2e^-16^. (G) Violin plot of RNA levels of the shared downregulated targets (*nosGAL4*>*stwl* GKD *& bamGAL4>stwl* GKD) in ovaries enriched for GSCs, cystoblasts, cysts, and whole ovaries, showing that Stwl downregulated targets are expressed at higher levels in the differentiated stages. Statistics: A negative binomial regression model was used to estimate the average TPM count of ‘signature’ genes between each ‘genotype’. The TPM of each gene was used as the dependent variable, and ‘genotype’ was the independent variable. Statistical comparisons between groups were performed using contrasts and p-values were adjusted for multiple comparisons using the Benjamini-Hochberg procedure (p-FDR). The average TPM between groups was considered to be significantly different when p-FDR < 0.05. (H) The biological process GO-terms of shared downregulated genes in ovaries of *nosGAL4*>*stwl* GKD *and bamGAL4>stwl* GKD compared with controls using fold enrichment. Statistics: Fisher’s exact test), showing that the overlapping downregulated targets are involved in egg formation as well as nucleosome assembly and DNA metabolic processes. Displaying results for FDR< 0.05.

We found 1496 genes (60%) of the upregulated genes are shared between *nosGAL4* and *bamGAL4-*mediated *stwl* GKD **(Figure 2B)**. Among these upregulated genes were *rpS19b* and *blanks,* consistent with our data showing that RpS19b::GFP and Blanks were ectopically expressed upon loss of *stwl* **(Figure 2A and 2C)**. We further validated that *rpS19b* RNA was upregulated in the *stwl* GKD egg chambers compared to the control by probing for its RNAs using *in situ* hybridization **(Figures S2A-S2B1)**. To determine if the other genes upregulated upon *stwl* GKD were early oogenesis genes, we plotted the abundance of the upregulated RNAs by using available RNA-seq libraries that were enriched for undifferentiated stages (GSCs and cystoblasts), differentiating stages when oocytes are specified (cysts), and differentiated stages (early egg chambers and late-stage egg chambers) **(Figure 2D)**.^16^ We found that the upregulated genes are expressed in the undifferentiated stages and then are repressed in the differentiated stages in a Stwl-dependent manner **(Figure 2D)**. GO-term analysis suggested that the genes upregulated upon loss of Stwl included cell-differentiation genes, consistent with them being early oogenesis genes **(Figure 2E)**. Some Stwl-regulated early oogenesis genes were also SETDB1-regulated early oogenesis genes, as we found a 48% overlap of the upregulated genes, including *rpS19b* and *blanks* **(Figure S2E)**.^15^ These data suggest that while Stwl and SETDB1 co-regulate the expression of genes, they also independently regulate a large cohort of genes. Thus, *stwl* promotes the silencing of genes, including a cohort of SETDB1-regulated early oogenesis genes, at the onset of oocyte specification.

Additionally, we found 370 genes (40%) of downregulated genes are shared between *nosGAL4* and *bamGAL4-*mediated *stwl* GKD **(Figure 2F)**. By plotting the abundance of the downregulated RNAs upon *stwl* GKD using available RNA-seq libraries that were enriched for different stages of oogenesis, we found that downregulated RNAs under control conditions were attenuated in the undifferentiated stages and are then expressed in the differentiated stages in an Stwl-dependent manner **(Figure 2G)**.^16^ The GO-term analysis of the downregulated genes suggested that the genes are involved in egg activation, meiosis I, and nucleosome assembly process, consistent with a failure in oogenesis and sterility upon loss of *stwl* **(Figure 2H)**. Downregulated genes included maternally deposited genes such as *polar granule component* (*pgc*) and *wispy* (*wisp*) **(Figure S2F)**.^63–65^ We validated that *pgc* RNA was downregulated in the *stwl* GKD egg chambers compared to the control by probing for its RNAs using *in situ* hybridization **(Figures S2C-2D1)**. However, other maternally provided RNAs, such as *hunchback,* were not downregulated **(Figure S2F)**.^66,67^ Thus, Stwl is required to upregulate a cohort of genes that are critical to promote oogenesis, fertility, and embryonic development.

### Stonewall binds at promoters and overlaps with boundary element proteins but does not directly regulate a large fraction of dysregulated genes

To determine whether Stwl regulates its targets directly, we conducted Cleavage Under Targets and Release Using Nuclease (CUT&RUN) using the Stwl antibody.^68,69^ We probed Stwl binding in the early stages of oogenesis in *bam* GKD ovaries, which are enriched for undifferentiated stages, and adult wild-type ovaries, which are enriched for differentiated egg chambers.^10,61^ We used *stwl* GKD ovaries and IgG as negative controls **(Figure 3A)**. We found Stwl binding to the genome increases from undifferentiated (CBs) to differentiated stages (egg chambers), consistent with the antibody staining showing an increase in Stwl levels in the cyst stages **(Figure S1H-S1I)**. The association of Stwl with the genome is reduced upon *stwl* GKD and are absent in the IgG-only sample **(Figure 3A)**. Taken together, we find that the Stwl binding signal from CUT&RUN during oogenesis is specific.

**Figure 3.**
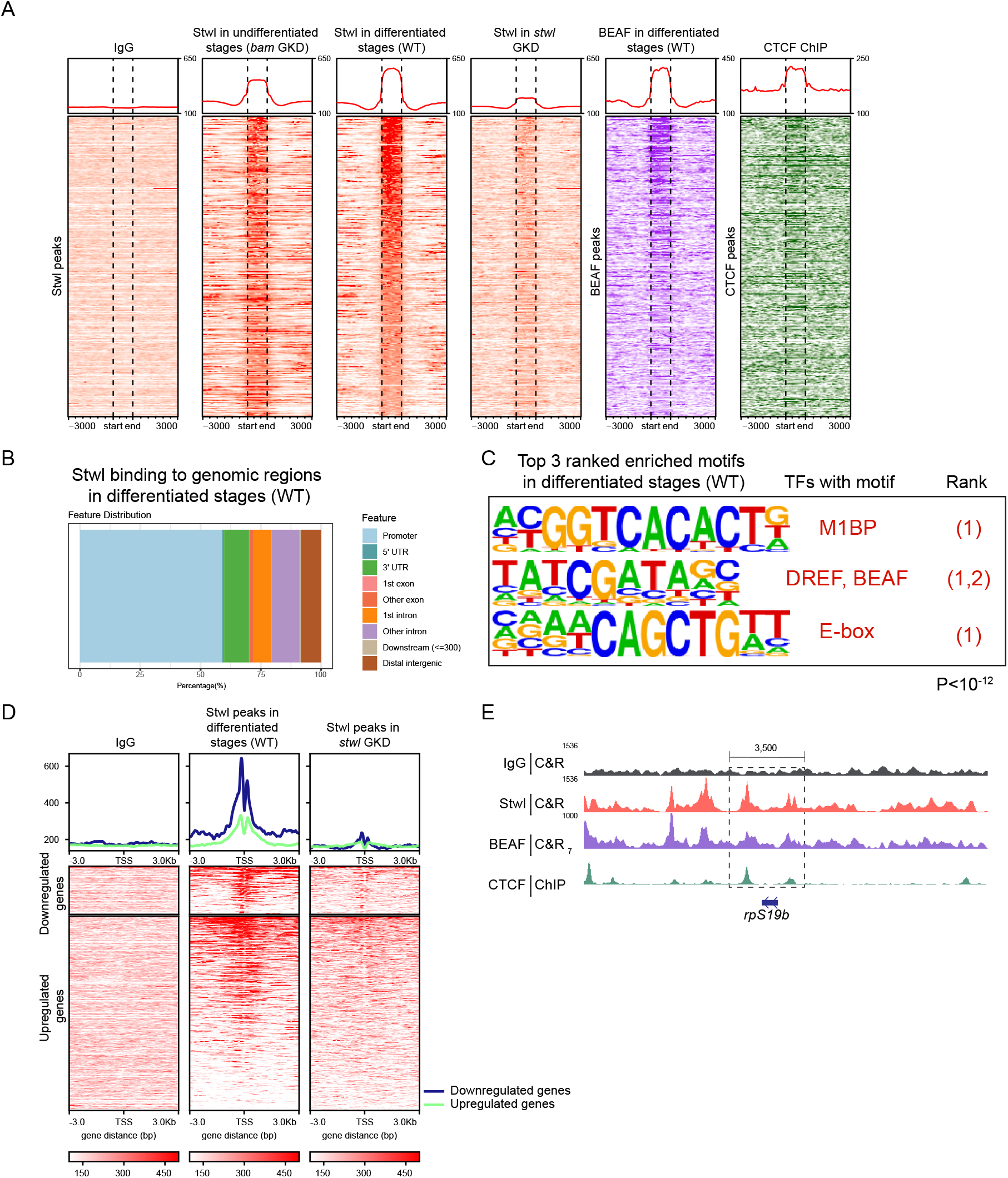
Stwl binds proximal to promoters and overlaps with boundary element proteins but does not directly regulate a large fraction of the dysregulated genes. (A) CUT & RUN occupancy of IgG peaks in ovaries (negative control), Stwl peaks in undifferentiated stages, Stwl peaks in differentiated stages (8585 peaks) as well as Stwl peaks upon *stwl* GKD, BEAF peaks in wild type (WT) differentiated stages (WT ovaries) in purple and CTCF ChIP-seq peaks in larval CNS in green. Heatmaps showing −3 Kb and +3Kb around the start and end of Stwl peaks shown in red. Black dashed lines represent the start and end of Stwl peaks. These heat maps show an overlap between Stwl, BEAF and CTCF binding. (B) Graph showing percentages of Stwl binding pattern to genomic regions in the differentiated stages. Light blue represents promoters, navy represents 5’UTRs, green represents 3’UTRs, salmon color represents 1st exons, red represents other exons, orange represents 1st introns, purple represents other introns, beige represents downstream (<300) and brown represents distal intergenic regions. This shows that Stwl binds mainly to promoters (60%). (C) Homer motif analysis of Stwl binding sites in the differentiated stages showing top 3 ranked motifs also bound by other transcription factors M1BP, DREF/BEAF, and E-box associated with a P<10^-12^. (D) CUT & RUN occupancy of IgG peaks in WT ovaries (negative control), Stwl peaks in differentiated stages (WT ovaries), as well as Stwl in *Stwl* GKD for RNA-seq upregulated and downregulated targets. Heatmaps showing −3 Kb and +3Kb around the TSS of target genes. These heat maps show Stwl, BEAF and CTCF bind strongly most downregulated targets and only a portion of the upregulated targets. (E) CUT & RUN occupancy track for *rpS19b* locus for IgG (negative control) shown in grey, Stwl peaks in differentiated stages of WT ovaries (red), BEAF peaks in differentiated stages of WT ovaries (purple) and CTCF ChIP-seq peaks in larval CNSs (green) showing Stwl, BEAF and CTCF flanking *rpS19b* locus from both sides.

In the differentiated egg chambers, Stwl was mostly bound at promoters and to a lesser extent in intergenic regions and introns **(Figure 3B)**. We performed motif enrichment analysis and found regions, including a TATCGATAGC motif, to be enriched **(Figure 3C)**. The enrichment of this motif, which is bound by proteins such as Boundary element-associated factor-32 (BEAF-32), is consistent with previous observations from ChIP-Seq carried out for Stwl in S2 cells **(Figure 3C)**.^51^ BEAF is an insulator protein that binds to boundary elements to regulate genome organization and chromatin state.^38,70,71^ To determine if Stwl is associated with boundary elements during oogenesis, we carried out CUT&RUN for BEAF and also used published ChIP-seq data for another boundary element protein CCCTC-Binding Factor (CTCF) **(Figure 3A)**.^72^ We found that Stwl binding regions partially overlapped with regions bound by BEAF and CTCF **(Figures 3A and S3A)**. Thus, Stwl is associated with promoter regions of the genome and overlaps with the insulator proteins BEAF and CTCF.

To determine if Stwl directly regulates early oogenesis genes, we integrated our RNA-seq and CUT&RUN data from control and *stwl* GKD ovaries. We found that in the differentiated stages, Stwl binds 23% of the upregulated targets and 44% of the downregulated targets above the baseline signal **(Figure 3D)**. Some upregulated early oogenesis genes, such as *rpS19b,* have Stwl binding sites while others, such as *blanks,* do not have any clear Stwl binding site proximal to the transcriptional start sites (TSS) **(Figures 3E and S3B)**. These data suggest that Stwl binds proximal to a cohort of upregulated and downregulated targets but does not directly regulate a large cohort of dysregulated genes.

### Stonewall is required to demarcate active and silenced genomic compartments during differentiation

Stwl exhibits distinct binding patterns: it binds at promoters of select upregulated targets and is also found at a large cohort of downregulated targets. This observation hints at a more intricate mechanism for how Stwl regulates gene expression beyond merely binding to targets and promoting their silencing. Further, we found that Stwl’s binding overlaps with insulator proteins BEAF and CTCF, which can demarcate genomic boundaries, including TADs.^36,39,73^ Based on these data, we hypothesized that Stwl may influence gene expression by influencing genomic boundaries during differentiation. The loss of Stwl could potentially affect the chromatin state and, consequently, gene expression. To test this hypothesis, we first probed Stwl’s binding patterns and its relationship with the chromatin state and genomic organization pre- and post-differentiation.

TADs exhibit remarkable conservation across different cell types.^74,75^ To determine where Stwl binds in relation to genome organization, we used previously annotated TADs from salivary gland cells and Kc167 cells to determine the binding sites of Stwl, CTCF, and BEAF in the context of genomic organization.^76–78^ We also conducted a comprehensive analysis of the chromatin state on control gonads by examining H3K4me3 (associated with active promoters), H3K27ac and H3K4me1 (associated with active enhancers), H3K36me3 (associated with transcribed gene bodies), and H3K27me3 and H3K9me3 (associated with gene silencing) chromatin marks using CUT & RUN pre- and post-differentiation.^79,79–82^ We used these chromatin marks to build a 7-state chromatin model **(Figure S4A)** and correlated TADs, Stwl/CTCF/BEAF binding sites, and the observed chromatin states.^83^ By exploring these interconnections, we aimed to gain a better understanding of the role of Stwl in regulating the overall chromatin landscape.

Although salivary gland cells had fewer annotated TADs than Kc167 cells, the salivary gland TADs were encompassed within the annotated TADs of Kc167 cells **(Figure S4B)**. In control ovarioles enriched for differentiated egg chambers, we found that Stwl, BEAF, and CTCF bound to regions flanking the regions annotated as TADs in Kc167 cells **(Figure 4A)**. We found that Stwl binding at the TAD boundaries increased from undifferentiated cells to differentiated egg chambers **(Figure 4A)**. *stwl* GKD results in loss of Stwl at these boundaries **(Figure 4A)**. As Stwl and BEAF binding overlap, we asked if Stwl could regulate BEAF at the boundaries. Through antibody staining, we found that BEAF expression was attenuated in the undifferentiated cells **(Figures S4C-S4C2)**. However, as the cysts differentiated into oocytes, we observed increased BEAF levels **(Figures S4C-S4C2)**. We found that loss of Stwl results in loss of BEAF at the boundaries **(Figure 4A)**. However, not all of BEAF’s genomic binding is subject to regulation by Stwl. We identified that BEAF also binds to genomic sites independently of Stwl, and in these cases, the loss of Stwl does not result in the absence of BEAF at these sites **(Figure S4D)**. Thus, Stwl and BEAF accumulate at TAD boundaries during GSC differentiation, and Stwl recruits BEAF to these boundaries either directly or indirectly.

**Figure 4.**
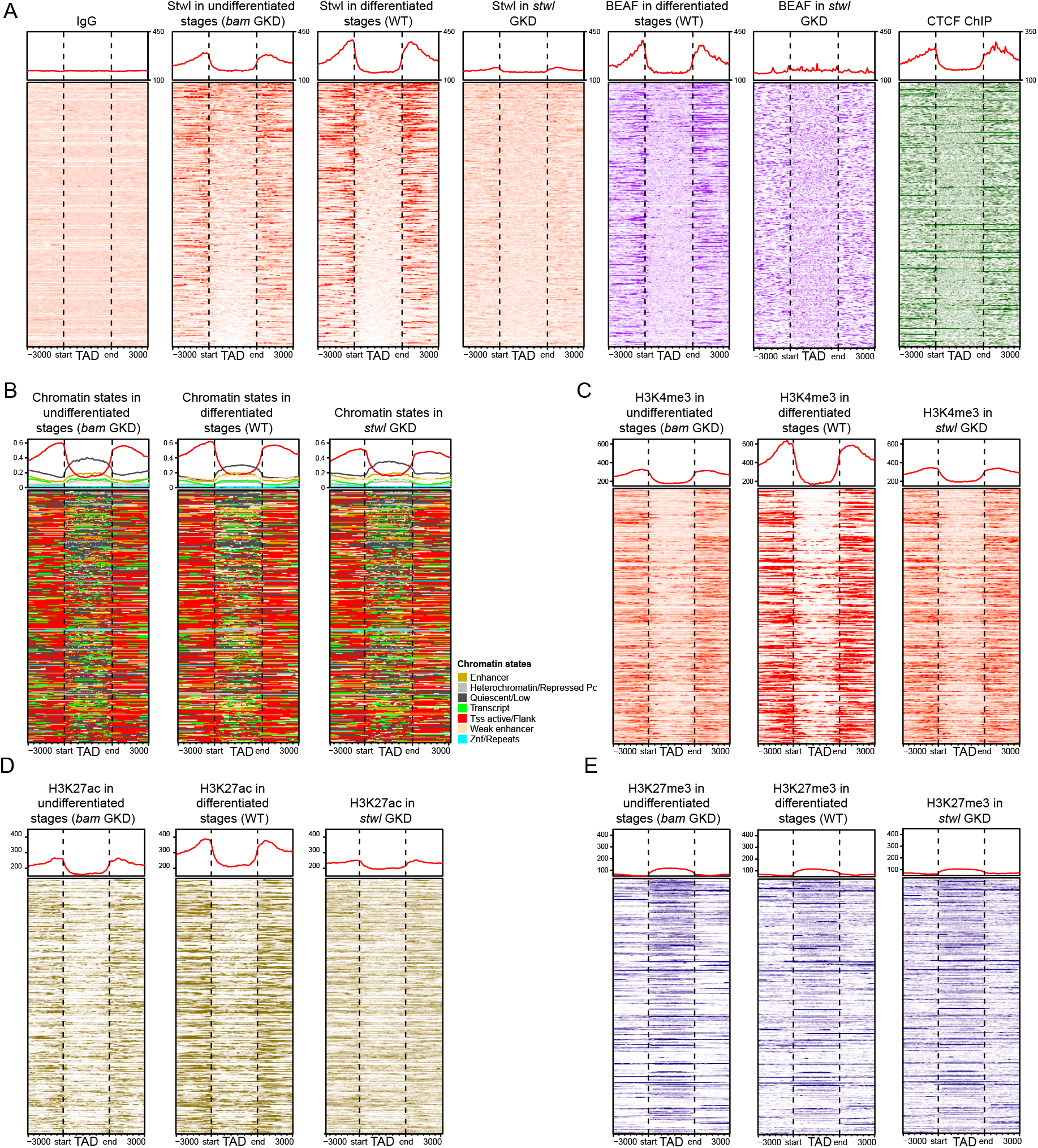
Stwl demarcates active and silenced genomic compartments during differentiation and is required for proper chromatin state. (A) CUT & RUN occupancy of IgG peaks in WT ovaries (negative control),Stwl peaks in undifferentiated stages (*bam* GKD), Stwl peaks in differentiated stages of WT ovaries as well as Stwl peaks in *stwl* GKD, BEAF peaks in differentiated stages of WT ovaries in *stwl* GKD and CTCF ChIP-seq peaks in larval CNSs. Occupancy heatmaps are shown for topologically associated domains (TADS) as well as −3Kb and +3 Kb around the start and end of TADs. Black dashed lines represent the start and end of TADs. The heatmaps show the binding of Stwl, BEAF and CTCF at the boundaries of TADS. (B) Comprehensive analysis of 7 chromatin states of undifferentiated stages (*bam* GKD), differentiated stages (WT ovaries), and *stwl* GKD ovaries built using multiple histone marks shown for TADs. Heatmap shown for TADS as well as −3Kb and +3 Kb around the start and end of TADs. Black dashed lines represent the start and end of TADs. Different states marked by colors (gold: enhancer, light grey: heterochromatin/repressed, dark grey: quiescent/low, green: transcript, red: TSS active/flank, beige: weak enhancer and blue: Znf/repeats). This 7 state model shows Stwl demarcates active compartments at TAD boundaries and repressed compartments in TADs. (C) Heatmaps of H3K4me3 active histone mark in undifferentiated stages (*bam* GKD), differentiated stages (WT ovaries) and *stwl* GKD ovaries shown for TADs. Heatmap shown for TADS as well as −3Kb and +3 Kb around the start and end of TADs. Black dashed lines represent the start and end of TADs. Heatmaps showing the H3K4me3 histone mark profile of *stwl* GKD resembles that of undifferentiated stages. (D) Heatmaps of H3K27ac active enhancer histone mark in undifferentiated stages (*bam* GKD), differentiated stages (WT ovaries) and *stwl* GKD ovaries shown for TADs. Heatmap shown for TADS as well as −3Kb and +3 Kb around the start and end of TADs. Black dashed lines represent the start and end of TADs. Heatmaps showing H3K27ac histone mark profile of *stwl* GKD resembles that of undifferentiated stages. (E) Heatmaps of H3K27me3 repressive histone mark in undifferentiated stages (*bam* GKD), differentiated stages (WT ovaries), and *stwl* GKD ovaries shown for TADs. Heatmap shown for TADS as well as −3Kb and +3 Kb around the start and end of TADs. Black dashed lines represent the start and end of TADs. Heatmaps showing H3K27me3 histone mark profile of *stwl* GKD resembling that of undifferentiated stages as well as differentiated stages.

We next examined the chromatin state of these compartments in the undifferentiated and differentiated stages, and *stwl* GKD. We found that in both the undifferentiated cells and differentiated egg chambers, Stwl binds at TAD boundaries and demarcates active chromatin (e.g. TSSs and enhancers) flanking the TADs from the quiescent genes present within the TADs **(Figure 4B)**. The increase in Stwl at these boundaries during differentiation coincided with changes to the chromatin state for both regions flanking and inside the TADs **(Figures 4B-D)**. For example, there was an increase in promoter activity (H3K4me3) in regions flanking the TADs during differentiation **(Figure 4C)**. There was also an increase in enhancer activity (H3K27ac) mostly at TADs boundaries and, at lower levels, within TADs in the differentiated stages **(Figure 4D)**. H3K27me3 within TADs did not appreciably change during differentiation **(Figure 4E)**. The changes associated with promoter and enhancer activity did not happen upon loss of *stwl* **(Figure 4A**, **4C-D)**. Thus, during differentiation, the binding of Stwl, BEAF, and CTCF demarcates active and silenced chromatin in TADs, which influences the chromatin state at TAD boundaries and within TADs.

### Stonewall regulates enhancer landscape to promote proper gene expression during differentiation

To determine how Stwl specifically affects gene expression, we examined the overlap between differentially expressed genes, TADs, and chromatin states in undifferentiated and differentiated controls as well as *stwl* GKD gonads. We found that the majority of upregulated genes (63%) and downregulated genes (56%) are encompassed within TADs **(Figure 5A)**. To determine how Stwl regulates upregulated and downregulated genes, we analyzed the chromatin state of these genes separately.

**Figure 5.**
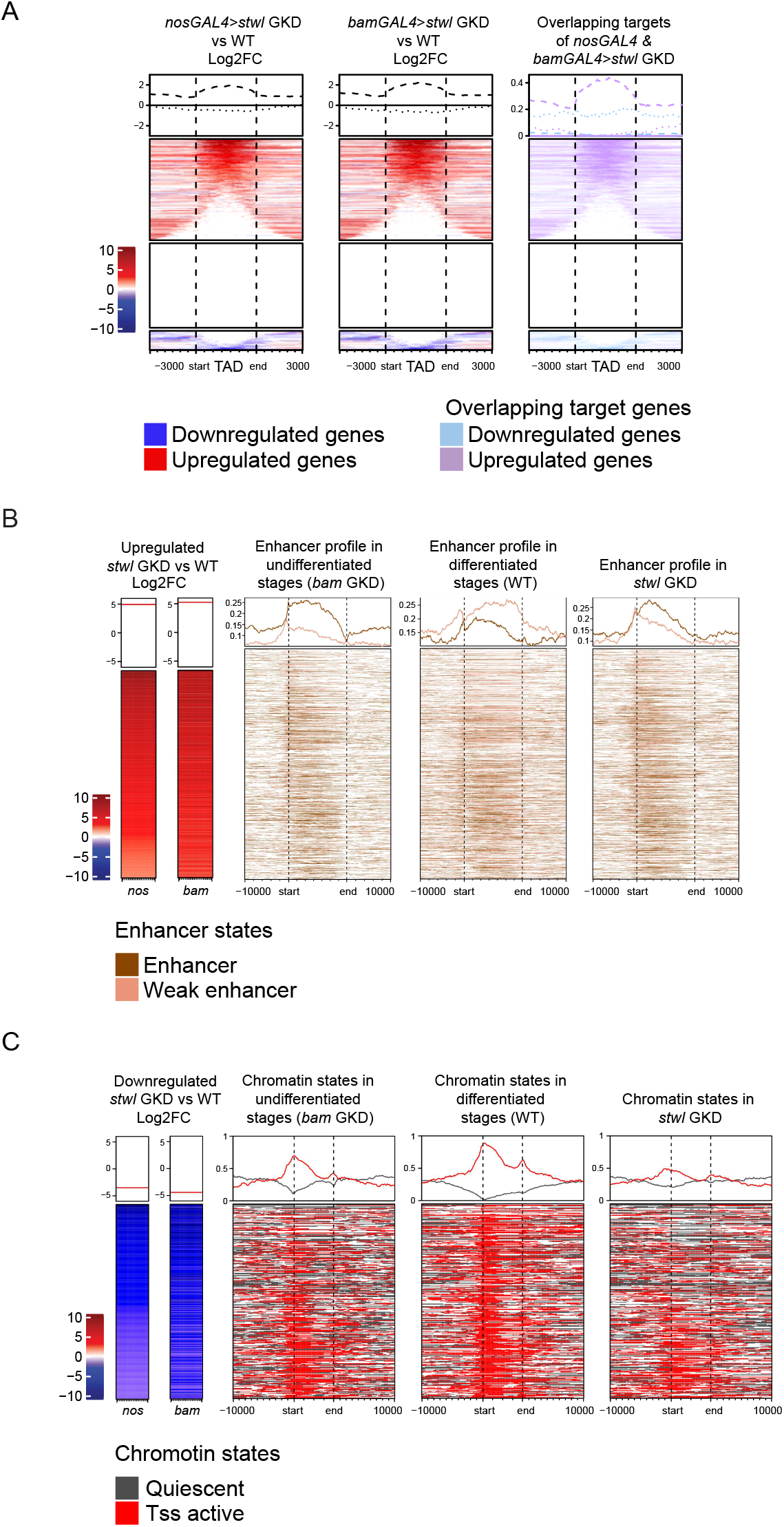
Stwl regulates enhancer activity to promote gene expression. (A) Heatmaps of lfc2 sorted for RNA-seq Stwl targets at TADs. Upregulated targets are shown in red and downregulated targets shown in blue. Overlapping targets shared between *nosGAL4>stwl* GKD and *bamGAL4>stwl* GKD showing upregulated targets in TADs (lilac) and downregulated targets (blue). Heatmaps shown for TADS as well as −3Kb and +3 Kb around the start and end of TADs. Black dashed lines represent the start and end of TADs. The heatmaps are divided into three clusters: the upregulated targets cluster, the cluster of non-target genes and the last cluster representing downregulated targets. The heatmaps show that both upregulated and downregulated genes upon *stwl* GKD are present within the genomic compartments. (B) Heatmaps of lfc2 sorted for RNA-seq Stwl upregulated targets in *stwl* GKD shown in red. Heatmaps show enhancers states for those targets in undifferentiated stages (*bam* GKD), differentiated stages of wild-type ovaries Enhancer profiles heatmaps are shown for upregulated targets in undifferentiated stages (*bam* GKD), differentiated stages of wild-type ovaries, and as well as *stwl* GKD ovaries showing changes in enhancer profiles during differentiation that do not occur upon *stwl* GKD. (C) Heatmaps of lfc2 sorted for RNA-seq Stwl downregulated targets in *stwl* GKD shown in blue. Heatmaps show chromatin states for those targets in undifferentiated stages (*bam* GKD), differentiated stages of wild-type ovaries, and as well as *stwl* GKD ovaries. Different states are marked by colors dark gray: quiescent/low, and red: TSS active/flank) showing an increase in active marks at TSS of genes and a decrease of quiescence during differentiation. These changes in chromatin states do not occur upon *stwl* GKD.

Analyzing the chromatin state of the upregulated genes upon *stwl* GKD, we observed changes in the enhancer landscape **(Figure 5B and S5A)**. In control ovaries, in the undifferentiated stages, genomic regions encompassing upregulated genes are enriched for active enhancers, consistent with the fact that these genes being normally expressed at higher levels in the undifferentiated stages **(Figures 5B and Figure 2D)**. During differentiation, we find that the levels of active enhancers decrease with an increase in “weak enhancers”, which are usually associated with inactive or poised genes, concomitant with these genes being silenced **(Figure 5B)**. These changes in “active” and “weak” enhancers during differentiation did not happen upon loss of *stwl* and these genes continued to be expressed **(Figure 5B and Figure 2D)**. The enhancer profile of *stwl* GKD closely resembles the enhancer profile of undifferentiated cells. Indeed, principal component analysis (PCA) plot of histone marks of upregulated genes shows that enhancer marks (H3K27ac) is dysregulated in *stwl* GKD compared to differentiated controls (**Figure S5C**). For example, the genomic region proximal to *rpS19b* showed a decrease of active enhancers and an increase of weak enhancers during differentiation **(Figure S5D)**. These changes in enhancer states are attenuated in *stwl* GKD ovaries **(Figure S5D)**. Thus, we infer that Stwl promotes silencing of a cohort of early oogenesis genes, such as *rpS19b.* during differentiation by regulating their promoter and their associated enhancers.

In contrast, in control ovaries during differentiation, the genes that are downregulated upon *stwl* GKD showed an increase in active marks at the TSS and a decrease in quiescence consistent with these genes being activated at this stage **(Figures 5C and S5B)**. However, upon *stwl* GKD, this increase in active marks does not occur at the promoters of the downregulated genes and these genes remain quiescent **(Figure 5C)**. The PCA plot shows that active marks such as H3K4me3 and H3K27ac are dysregulated in *stwl* GKD compared to differentiated controls. The chromatin state of the promoters of the downregulated genes upon *stwl* GKD is closer to the undifferentiated stages than the differentiated stages **(Figure S5C)**. For example, *pgc* and *wispy*, downregulated genes, show an increase in active marks during differentiation and a decrease in quiescence. The decrease in quiescence and increase in the active marks is attenuated upon *stwl* GKD **(Figure S5E)**. Thus, Stwl contributes to expression of a cohort of maternal genes by promoting the acquisition of active histone marks during differentiation.

### Stonewall promotes the expression of Nucleoporins and thus NPC formation and the anchoring of silenced genes to the lamina during differentiation

We were interested in understanding how Stwl, which is present at TAD boundaries, contributes to changes to the expression of genes inside TADs. NPCs can help position TADs to the nuclear lamina to maintain their silenced state.^44–46,84–86^ We previously described a role for NPCs in maintaining the silencing of early oogenesis genes such as *rpS19b* by tethering them to the nuclear periphery.^15^ Intriguingly, loss of individual Nups in the germline phenocopies loss of *stwl*.^15,23^ Building upon these findings, we asked if Stwl affects the expression of Nups.

From visual inspection of downregulated targets, we observed a significant decrease in the expression levels of 4 nucleoporins (Nups), which were downregulated by greater than 2-fold upon the loss of *stwl* **(Table 1**, **Figure 6A)**. Another 10 were also downregulated but did not meet the 2-fold cutoff. These downregulated Nups did not belong to a specific NPC subcomplex. Instead, we discovered that these Nup genomic loci were bound by Stwl and BEAF around the TSS (**Figure 6A)**. For example, for the Nup *aladin,* which is downregulated upon the loss of *stwl*, there is a decrease of quiescence in the genomic region proximal to it during differentiation. This change in quiescence did not occur upon *stwl* GKD **(Figure S6A)**.^87^ Taken together, we find that Stwl promotes the expression of a specific cohort of Nups, by creating a barrier between active and repressed chromatin regions during differentiation.

**Figure 6.**
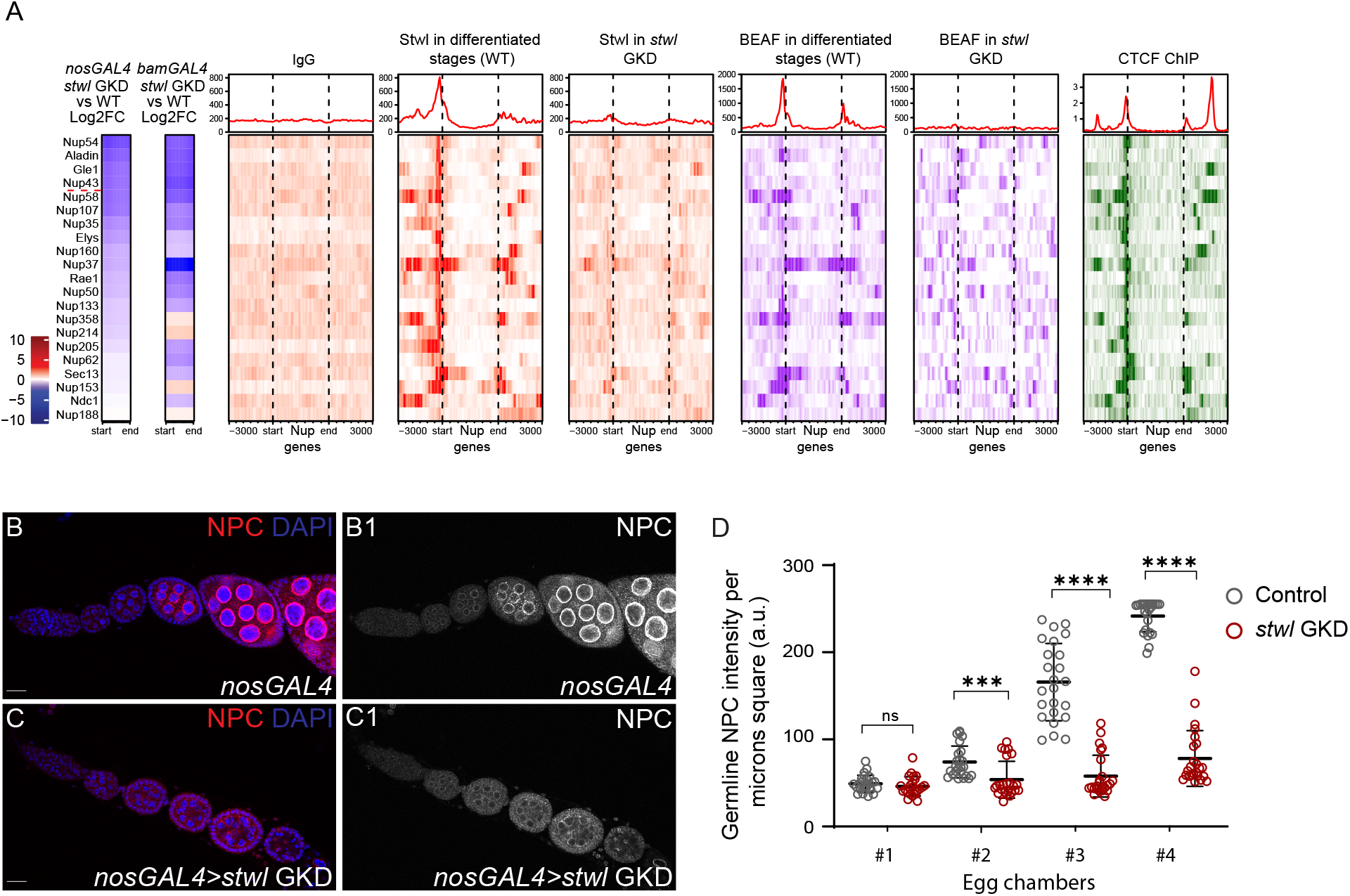
Stwl promotes the expression of Nucleoporins to promote NPC formation. (A) CUT & RUN occupancy of IgG peaks in WT ovaries (negative control), Stwl peaks in undifferentiated stages of ovaries, Stwl peaks in differentiated stages of WT ovaries as well as Stwl peaks upon *stw*l GKD as well as BEAF peaks in differentiated stages of WT and CTCF ChIP-seq peaks in larval CNSs for Nucleopore complex genes (Nups). Occupancy heatmaps are shown for Nups genes as well as −3Kb and +3 Kb around the start and end of Nups genes.Black dashed lines represent the start and end of Nups.The heatmaps show the binding of Stwl, BEAF, and CTCF binding of Nucleopore complex genes (Nups).The red dashed line delineates the Nups that are downregulated by lfc2.Heatmaps show binding of Stwl, BEAF and CTCF at TSS of *nup* genes. (B-B1) Ovariole of control *nosGAL4* (B) and grayscale (B1) stained for NPC (red, right grayscale) and DAPI (blue). NPCs are forming regular ring structures and increase in intensity, leading to the later egg chambers. (C-C1) Ovariole of *stwl* GKD (C) and grayscale (C1) stained for NPC (red, right grayscale) and DAPI (blue). NPCs do not increase in intensity leading to the later egg chambers. (D) Arbitrary unit (a.u.) quantification of nucleopore genes (Nups) in (2×2) microns square of egg chambers of *stwl* GKD (red) compared with control ovaries (grey). In the control, NPCs are expressed in egg chambers, and their intensity per micron square increases throughout development. In *stwl* GKD, NPCs intensity is not increasing.Statistics: Two tailed t-test; n = 5 ovarioles per genotype; ns, p > 0.05; ∗p < 0.05; ∗∗p < 0.01; ∗∗∗p < 0.001; ∗∗∗∗p < 0.0001.

Considering that all Nups are required to form functional NPCs, we hypothesized that the loss of *stwl* would disrupt NPC assembly. To test this hypothesis, we examined NPC formation in control and *stwl* GKD conditions utilizing the mAb414 antibody, a marker for NPCs.^15,21,43,88^ From immunostaining, we found that NPC levels did not increase in egg chambers in *stwl* GKD ovaries as they did in the controls **(Figures 6B-6D).** In addition, when there was NPC staining present in *stwl* GKD ovaries, we found that there were gaps in the distribution of NPCs in the nuclear membrane **(Figures S6B-S6C1)**. Thus, Stwl is required for proper NPC formation.

Chavan *et al.*, have shown that Stwl physically interacts with components of NPC. We hypothesized that Stwl, via interaction with NPCs, can promote the association of TADs to the lamina.^44,84,89^ To assess this, we analyzed published Lamin C ChIP data from S2 cells and discovered that Stwl binding sites overlap with, and are proximal to Lamin C binding sites, including at TAD boundaries **(Figure 7A-7B)**.^89^ We examined the TAD that encompasses the early oogenesis gene *blanks* and found that not only was the TAD boundaries associated with Lamin C, the *blanks* locus was also associated with Lamin C **(Figure S7A)**. We also examined the early oogenesis gene *rpS19b* which is not part of a TAD and found that Stwl binding sites that flank this locus overlap with Lamin C **(Figure S7A)**. To further support the hypothesis that Stwl promotes tethering of silenced genes to the lamina, we performed DNA fluorescent *in situ* hybridization (FISH) using probes for the *rpS19b* locus in control and *stwl* GKD ovaries.^15^ In control ovarioles, we observed that the *rpS19b* locus was located at the nuclear lamina, consistent with previous observations **(Figure 7C)**. However, upon *stwl* GKD, the *rpS19b* locus was no longer positioned at the nuclear periphery, indicating the untethering of this locus from the lamina **(Figures 7C-7E)**. Thus, Stwl promotes the association of silenced genes with the nuclear lamina to help maintain the silenced state of genes.

**Figure 7.**
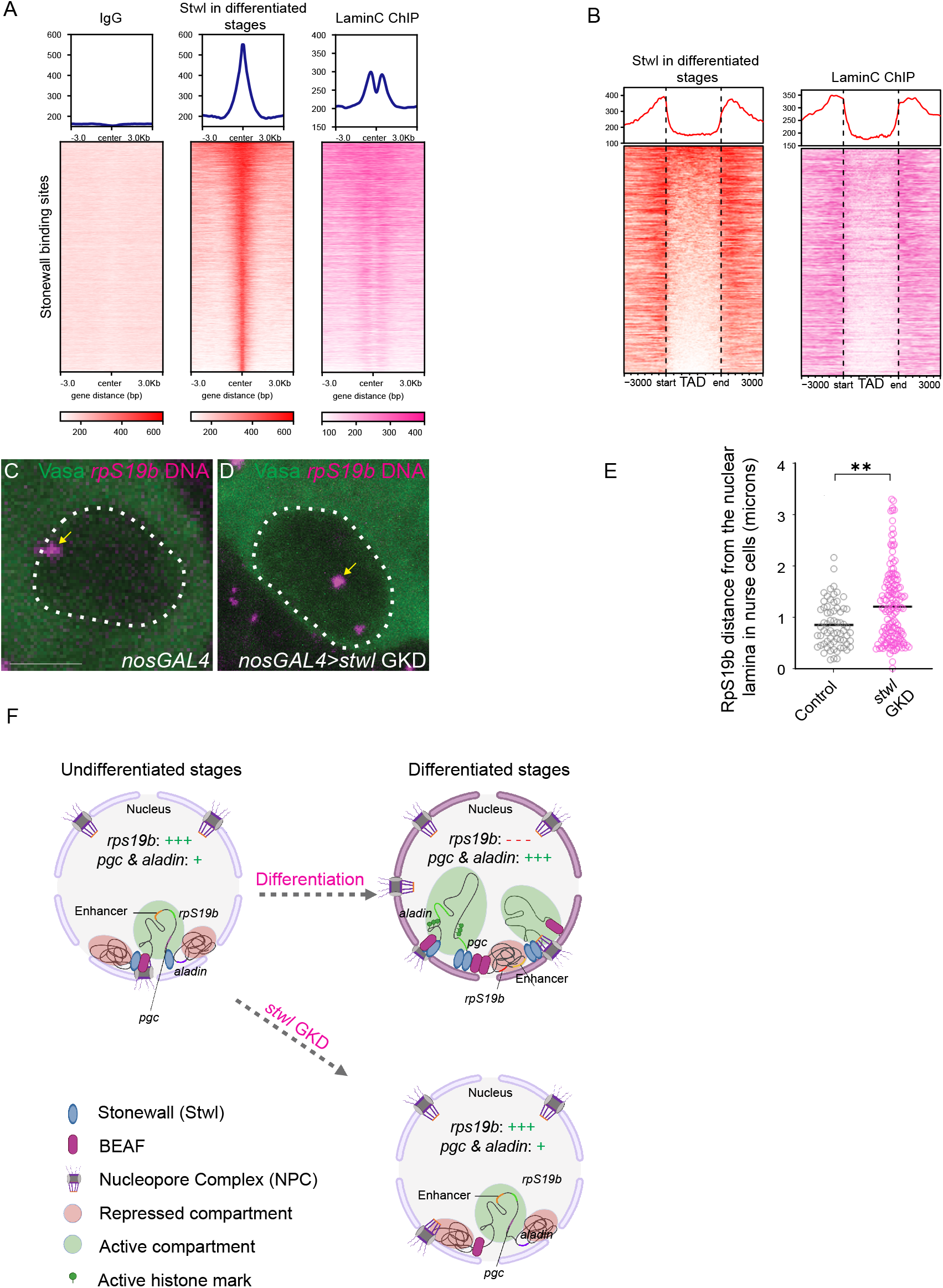
Stwl promotes association of genes to the nuclear periphery. (A) CUT&RUN occupancy of Stwl and Lamin C peaks in differentiated stages of WT ovaries for all stwl binding sites. Stwl peaks in differentiated stages are shown in red and Lamin C peaks from Lamin C ChIP in S2 cells shown in pink. Heatmaps showing Stwl and Lamin C peaks Stwl binding sites as well as −3Kb and +3 Kb around. Lamin C is overlapping and proximal to Stwl binding sites. (B) CUT&RUN occupancy of Stwl and Lamin C peaks in differentiated stages of WT ovaries at TADs. Stwl peaks are shown in red and Lamin C peaks in pink. Black vertical dashed lines represent the start and end of TADs as well as −3Kb and +3 Kb around them. Lamin C is observed to be enriched proximal and at Stwl binding sites at TAD boundaries. (C-D) Nurse cell nucleus of control (C) and *stwl* GKD ovaries (D) stained for Vasa (green) and probed for *rpS19b* DNA (magenta). In control, *rpS19b* is at the nuclear periphery*. stwl* GKD resulted in the migration of *rpS19b* away from the nuclear lamina that was tethering it. (E) *rpS19b* distance measurement from the nuclear lamina in nurse cells of control WT ovaries and *stwl* GKD ovaries. Distance is measured in microns, shown in grey for control and pink for *stwl* GKD. In control, RpS19b is close to the nuclear lamina in contrast to in *stwl* GKD where they are further from the nuclear lamina.Statistics: unpaired t-test; n =4 ovarioles per control and n=8 per *stwl* GKD; ns, p > 0.05; ∗p < 0.05; ∗∗p < 0.01; ∗∗∗p < 0.001; ∗∗∗∗p < 0.0001. (F) Model: Stwl is required during the transition from undifferentiated stages to differentiated stages of oogenesis to establish boundaries between active and repressed domains by recruiting BEAF to its binding sites and regulating the expression of germ cell differentiation genes and Nups through enhancer dynamics at the nuclear lamina. This model was created with BioRender.com. Scale bars: 15 μm

## Discussion

Germ cell differentiation into an oocyte involves significant changes in gene expression.^15,56,57,90–92^ Germ cell-specific genes are silenced during this transition, while maternally deposited genes are activated.^15,92^ Active genes tend to be in the nuclear interior, whereas inactive genes are mainly found near the nuclear periphery, close to the lamina.^44,84^ The mechanisms that promote such genomic organization during the germ cell to maternal transition had not been deciphered. Here, we found that Stwl is present at TAD boundaries delineating active and repressed genomic compartments. The presence of Stwl at these boundaries facilitates the establishment of specific chromatin states of these genomic compartments and the maintenance of chromatin marks. Demarcating these compartments is required for both silencing germ cell genes and activating a cohort of maternal genes. In addition, the Stwl-dependent formation of genomic boundaries also promotes the expression of Nups, which aid in the formation of NPCs **(Figure 7F)**.^22,93^ These NPCs, in turn, assist in tethering repressed regions to the nuclear lamina. Thus, Stwl regulates local and global genome organization to regulate the cell fate transition during oogenesis **(Figure 7F).**

### Stonewall silences early oogenesis genes by demarcating genomic compartments

Stwl is required for GSC maintenance and the proper development of egg chambers.^52^ Previously, it had been proposed that Stwl silences differentiation genes potentially via epigenetic control by modulating H3K9me3 and H3K27me3.^51,94^ Indeed, we find that Stwl is critical for silencing early oogenesis genes, some of which promote GSC differentiation into an oocyte.^95^ However, we believe that this function of Stwl is not direct. Instead, Stwl plays a crucial role in demarcating silenced and active compartments within the genome. By regulating the distribution of these chromatin marks, Stwl indirectly promotes the silencing of early oogenesis genes during oogenesis. There seem to be two classes of genes that Stwl regulates. One class of genes, such as *rpS19b*, are not in TADS but contain H3K9me3 on their gene bodies^15^, and are flanked by Stwl/BEAF binding sites. The other class of genes such as *blanks* are not proximal to Stwl binding and instead are inside TADs that are quiescent. Loss of Stwl from the boundaries of these TADs leads to the changes to chromatin state inside TADs and appearance of an enhancer proximal to *blanks*. Thus, Stwl can act as a barrier at the level of individual genes and genomic compartments to promote the silencing of early oogenesis genes.

The genomic compartments that Stwl demarcates during oogenesis are annotated as TADs in Kc167 cells and salivary gland cells.^77,78^ The function of TADs has been shown to constrain promoter-enhancer interactions to prevent enhancer capture to regulate proper gene expression.^39,41,42^ It is possible that ectopic expression of repressed genes, such as early oogenesis genes, due to loss of *stwl* could be because of enhancer capture caused by loss of partition between active and repressed chromatin. However, we do not know if the genomic compartments we have identified during oogenesis are bonafide TADs during oogenesis.

Stwl is not only required for silencing gene expression but is also critical for activating a cohort of maternally supplied genes such as *pgc* and some *Nups*.^96,97^ We do not think that these maternally provided genes are directly activated by Stwl. Instead, the maternal genes regulated by Stwl are proximal to genomic boundaries that separate active and repressed regions. We think a cohort of maternally supplied genes requires the activity of Stwl to provide a barrier from silenced regions present proximally. While the cohort of maternal genes regulated by Stwl is small compared to the number of genes supplied maternally, they are functionally critical. For example, *pgc* is required to specify a germ cell fate, and Nups are required to complete oogenesis and launch the next generation.^15,96^ Thus, Stwl coordinates stable silencing of early oogenesis genes to activation of maternally provided genes by providing a barrier function between chromatin compartments, preventing the mixing of chromatin states.

### Stonewall coordinates global genome reorganization and NPC formation

NPCs are required for global genome organization, but whether global genome organization itself promotes NPC formation was not known.^21,22^ Here, we found that Stwl, by acting as a barrier, between active and repressive compartments, promotes the expression of a cohort of Nups. As all Nups are required for NPC formation, Stwl promotes NPC formation by promoting the transcription of Nups that comprise the NPC. We previously showed that H3K9me3-mediated chromatin marks are also required for Nup transcription.^15^ This suggests that silencing marks and the ability to partition these silencing marks are critical for NPC formation.

NPCs are known to help in a genomic organization by helping tether both silenced genes to the lamina to maintain their silenced state and position active genes under NPCs to allow for RNA export.^86,97,98^ We find that loss of *stwl* results in loss of lamin association of an early oogenesis gene *rpS19b*. Chavan *et al.* show that Stwl also physically interacts with NPCs, promoting the anchoring of silenced genes to the nuclear periphery. Taken together, the data suggest that Stwl regulates NPC formation and then, by interaction with NPCs, anchors TADs to the nuclear periphery to maintain their silencing. Thus, Stwl modulates global genome organization by regulating the formation of and binding to NPCs.

Taken together, we find that Stwl is a critical regulator of local and global genome architecture during the germ cell to maternal transition. By establishing boundaries between silenced and active regions, Stwl ensures the confinement of a particular chromatin state and the proper expression of germ cell differentiation genes and Nups to regulate NPC formation. The NPCs, in turn, promote the tethering of silenced regions to the lamina. This work provides an essential framework for understanding the interplay between genome organization and cell fate determination.

## Materials and Methods

### Fly lines

The following RNAi and mutant fly stocks that were used in this study are: *Stwl* RNAi (Bloomington #35415), *stwl* deficiency chromosome (*Df(3L)Exel6122*) (Bloomington #7601), *Stwl* mutants using CRISPR by precise deletions of the open reading sequence obtained from the Jagannathan lab, *Nup54* RNAi (Bloomington # 57426).

The following tagged lines were used in this study: *rpS19b-GFP* (Buszczak Lab)^16^.

The germline-specific drivers and double balancer lines that were used in this study are: *UAS-Dcr2;nosGAL4* (Bloomington #25751), *bamGAL4* (Bloomington #80579),*matGAL4* (Bloomington #7062,7063),*nosGAL4;MKRS*/TM6 (Bloomington #4442), and *If*/CyO*;nosGAL4* (Lehmann Lab).

### Reagents for fly husbandry

Fly crosses were grown at 25-29°C and dissected between 0-3 days post-eclosion. Fly food for stocks and crosses were prepared using the Lehman lab protocol (summer/winter mix) and narrow vials (Fisherbrand Drosophila Vials; Fisher Scientific) were filled to approximately 10-12mL.

### Dissection and Immunostaining

Ovaries were dissected and the ovarioles were separated using mounting needles in PBS solution and kept on ice. Samples were then fixed for 12 minutes in 5% methanol-free formaldehyde. Ovaries were washed in 0.5 mL PBT (1X PBS, 0.5% Triton X-100, 0.3% BSA) 4 times for 10 minutes each while incubating on a nutator. Primary antibodies in PBT were added and incubated at 4°C nutating overnight. Samples were next washed 3 times for 5-8 minutes each in 1 mL PBT. Secondary antibodies were added in PBT with 4% donkey serum and incubated at room temperature for 3-4 hours. Samples were washed 3 times for 10 minutes each in 1 mL of 1X PBST (0.2% Tween 20 in 1x PBS) and incubated in Vectashield with DAPI (Vector Laboratories) for at least 1 hour before mounting.

The primary antibodies used are: rabbit anti-Stwl 1 (1:2000, obtained from the Jagannathan Lab) mouse anti-1B1 (1:20; DSHB), Rabbit anti-Vasa (1:1,000; Rangan Lab), Chicken anti-Vasa (1:1,000; Rangan Lab), Rabbit anti-GFP (1:2,000; abcam, ab6556), Rabbit anti-H3K9me3 (1:500; Active Motif, AB_2532132), Mouse anti-H3K27me3 (1:500; abcam, ab6002), Rabbit anti-Egl (1:1,000; Lehmann Lab), Mouse anti-NPC (1:2000; BioLegend, AB_2565026). The following secondary antibodies were used: Alexa 488 (Molecular Probes), Cy3 and Cy5 (Jackson Labs) were used at a dilution of 1:500.

### Fluorescence Imaging

Ovaries were mounted on slides and imaged using a Zeiss LSM-710 and LSM-880 confocal microscope under 20X, 40X and 63X oil objective with pinhole set to 1 airy unit. Image processing was done using Fiji and gain adjustment and cropping were performed in Photoshop.

### RNA isolation & TURBO

Ovaries were dissected into PBS and transferred to RNase free microcentrifuge tubes. PBS was removed and 100ul of Trizol was added and ovaries were flash frozen and stored at −80 C. Ovaries were then lysed in the microcentrifuge tube using a plastic disposable pestle. Trizol was added to 1 mL total volume while vigorously shaking the tubes and incubated for 5 min at RT. The samples were centrifuged for 20 min at >13,000 g at 4 C, and the supernatant was transferred to a fresh microcentrifuge tube. 500 ul of chloroform was added, and the samples were vigorously shaken and incubated for 5 minutes at RT. Samples were spun at max speed for 10 minutes at 4 C. The supernatant was transferred to a fresh microcentrifuge tube and ethanol precipitated. Sodium acetate was added, equaling 10% of the volume transferred and 2-2.5 volumes of 100% ethanol were added. The samples were shaken thoroughly and left to precipitate at −20 C overnight. The samples were centrifuged at max speed at 4 C for 15 min to pellet the RNA. The supernatant was discarded and 500 ul of 75% ethanol was added to wash the pellet. The samples were vortexed to dislodge the pellet to ensure thorough washing. The samples were spun at 4 C for 5 min and the supernatant was discarded. The pellets were dried for 10-20 min and then resuspended in 20-50ul of RNAse-free water and the absorbance at 260 was measured on a nanodrop to measure the concentration of each sample.

### RNA-seq library preparation and analysis

Libraries were prepared using the Biooscientific kit (Bioo Scientific Corp., NOVA-5138-08) and following their protocol. RNA was prepared with Turbo DNAse according to the manufacturer’s instructions (TURBO DNAfree Kit, Life Technologies, AM1907), and incubated at 37C for 30 min. DNAse was inactivated using the included DNAse Inactivation reagent and buffer according to the manufacturer’s instructions. The RNA was centrifuged at 1000 g for 1.5 min and 19 ml of supernatant was transferred into a new 1.5 mL tube. This tube was again centrifuged at 1000 g for 1.5 min, and 18 ml of supernatant was transferred to a new tube to minimize any inactivation reagent carry-over. RNA concentration was measured on a nanodrop. Poly-A selection was performed on a normalized quantity of RNA dependent on the lowest amount of RNA in a sample. Poly-A selection was performed according to the manufacturer’s instructions (Bioo Scientific Corp., 710 NOVA-512991). Following Poly-A selection mRNA libraries were. Library quantity was assessed via Qubit according to manufacturer’s instructions and library quality was assessed with a Bioanalyzer or Fragment Analyzer according to manufacturer’s instructions to assess the library size distribution. Sequencing was performed on biological duplicates from each genotype on an Illumina NextSeq500 by the Center for Functional Genomics (CFG) to generate single-end 75 base pair reads. Libraries were first evaluated for their quality using FastQC (v0.11.8, RRID:SCR_014583).^99^ Trim Galore! (v0.6.6, RRID:SCR_011847) was used to trim the adapter sequences with a quality threshold of 20.^100^ Reads were aligned to the dm6 reference genome version for drosophila melanogaster using STAR aligner (v2.7.5b, RRID:SCR_004463).^101^ Gene level read counts are obtained by using Salmon (v1.2.1, RRID:SCR_017036) for all libraries.^102^ Sample normalization was carried out using the median-ratios normalization method from DESeq2 R package (v1.30.1, RRID:SCR_015687), and differential expression analysis was performed using DESeq2.^103^ Genes with less than 5 reads in total across all samples are filtered as inactive genes. A gene is considered differentially expressed if the Benjamini-Hochberg adjusted p-value is less than 0.05 and the absolute log2 fold change is greater than or equal to 2. Volcano and violin plots were generated by ggplot2^104,105^ R (v.4.3.1) was used to perform the analysis.

### *In situ* hybridization

The *in-situ* hybridization procedure for *Drosophila* ovaries was followed.^15,17^ Probes were designed and generated through LGC Biosearch Technologies using Stellaris® mRNA FISH Probe Designer. Ovaries (3 pairs per sample) were dissected in RNase-free 1X PBS and fixed in 1 mL of 5% formaldehyde for 10 minutes. The ovaries were then permeabilized in 1mL of permeabilization Solution (PBST+1% Triton-X) while nutating at RT for 1 hour. Samples were then washed in a wash buffer for 5 minutes (10% deionized formamide and 10% 20x SSC in RNase-free water). Ovaries were covered and incubated overnight with 1ul of probe in hybridization solution (10% dextran sulfate, 1 mg/ml yeast tRNA, 2 mM RNaseOUT, 0.02 mg/ml BSA, 5x SSC, 10% deionized formamide, and RNase-free water) at 30°C. Samples were then washed 2 times in 1 mL wash buffer for 30 minutes and mounted in Vectashield.

### Hybridization Chain Reaction (HCR) for Drosophila ovaries

This protocol was adapted from Slaidina and Lehmann et al. 2021 and Molecular Instruments’ sample-in-solution protocol.^56^ All experimental steps were conducted using RNase-free reagents and equipment, including conical tubes, eppendorf tubes, and PCR tubes. For processes requiring a temperature of 37°C, samples were placed on a heat block. On the first day,Drosophila ovaries tissues were dissected in RNase-free 1X phosphate-buffered saline (PBS). Next, a fixation step involved the addition of 500 uL of 10% paraformaldehyde (PFA) to 750 uL of a solution containing 0.1% Tween-20 in PBS, and the samples were fixed for 20 minutes at room temperature (RT). Subsequently, the samples were washed twice with 500 uL of 0.1% Tween-20 in PBS for 5 minutes at RT. The samples were dehydrated using a series of dilution of absolute ethanol solutions, and then stored at −20°C for a minimum of overnight, up to 7 days. On the second day, the samples underwent rehydration, transitioning through a series of ethanol solutions on ice. Then, samples were permeabilized with 1% Triton X-100 in PBS at RT for 2 hours. Post-fixation was achieved by adding 500 uL of 10% PFA to 750 uL of 0.1% Tween-20 in PBS for 20 minutes at RT, followed by two 5-minute washes with 0.1% Tween-20 in PBS on ice. Subsequent washes involved 5xSSCT for 5 minutes on ice, and the samples were incubated in probe hybridization buffer for 5 minutes on ice, with pre-warming of the buffer at 37°C. Pre-hybridization was carried out in the pre-warmed probe hybridization buffer for 30 minutes at 37°C. The probe solution, consisting of 16 uL of probes in 1 mL of pre-warmed probe hybridization solution, was prepared and used for hybridization at 37°C overnight. On the third day, the samples were washed four times with warmed probe wash buffer for 15 minutes at 37°C and three washes with 5xSSCT for 5 minutes at RT. Amplification buffer, pre-warmed to RT, was then incubated with the samples for 30 minutes. Hairpin solutions for each specific amplifier probe were prepared by separately mixing 10 uL of 3uM stock of each hairpin 1 and hairpin 2. The resulting mixture was snap-cooled and added to 500 uL of amplification buffer at RT. Samples were incubated in the hairpin solution overnight at RT. Notably, all RT steps should be performed on a heat block set to 25°C.The final day involved a series of washes with 500 uL of 5xSSCT at RT. These included two 5-minute washes, two 30-minute washes, and one 5-minute wash. Finally, the samples were stored in Prolong mounting media with DAPI at 4°C in the dark for at least overnight.

### DNA FISH with IF

The DNA FISH procedure from the Jaganathan lab was followed. 3-4 ovaries were dissected in PBS and teased. Then, they were fixed in the fixative solution (50ul 16% formaldehyde,1ul NP40,20ul 10X PBS, 429ul water) for 10 minutes. The ovaries were then washed 3 times in PBX (50 ml 1X PBX and 150ul Triton-X) for 5 minutes each. The sample was then incubated in a primary antibody and PBX mixture overnight at 4C. The sample was washed 3 times (1 quick and 2 5-minute washes) in PBX the next day. Secondary antibodies were then added to the sample in PBX and incubated for 2 hours at room temp. Following this, the sample was washed first in 2X SSCT, then in 20% formamide-SSCT for 10 minutes, followed by another 10-minute wash in 40% formamide-SSCT and one last one in 50% formamide-SSCT. The ovaries were then transferred to a tube in 50% formamide-SSCT and pre-denature the sample 37C for 4 hours, 92C for 3 minutes, and 60C for 20 minutes. The samples were then transferred into a fresh tube, the formamide solution was removed, and a 37ul probe buffer (50 ul formamide,50ul 4X SSCT, 100ul 20% dextran sulfate) + 1ul RNase A were added. Following this, 3ul of the probe was added and mixed by pipetting. The sample was then incubated at 91C for 3 minutes then at 37 C overnight. The sample was washed twice with 50% formamide-SSCT for 30 minutes at 37 C while shaking, followed by a minute wash for 5 minutes at room temp in 20% formamide-SSCT. Following this, one last quick wash with PBS was performed, and Vectashield was added, followed by sample mounting.

### CUT & RUN assay

Before starting the experiment, stock solutions were prepared and stored: Wash Buffer (47.5 mL H2O 1 mL 1 M HEPES pH7.5 final 20 mM 1.5 mL 5 M NaCl final 150 mM 50 mg BSA final 0.1%), 1X BBT (0.5 g BSA final 0.5% 50 ml PBST), 2X STOP buffer (46 mL H2O 2 mL 5 M NaCl final 200 mM 2 mL 0.5 M EDTA final 20 mM), 100 mM CaCl2 solution, MXP buffer (10 g PEG8000 final 20% 25 mL 5 M NaCl final 2.5 M 0.5 mL 1 M MgCl2 final 10 mM Fill up to 50 mL with H2O), and permeabilization buffer (50mL PBST 500 μL Triton-X). On the first experimental day, the wash buffer+ and the BBT+ were freshly prepared and lasted up to 3 days. The wash buffer+ consisted of adding one large Roche complete EDTA-free tablet and 5 μL of 5.55 M Spermidine to achieve a final concentration of 0.5 mM and the BBT+ consisted of adding a large Roche complete EDTA-free tablet along with 5 μL of 5.55 M Spermidine to attain a final concentration of 0.5 mM and 200 μL of 0.5 M EDTA for a final concentration of 2Mm. Following the preparation, 20 pairs of fattened fly ovaries were dissected per replicate and placed on ice in 1X PBS. The sample was then treated with the permeabilization buffer for 1 hour at room temperature while nutating, followed by washing with 1 mL BBT+ buffer and subsequent removal of the supernatant. Antibody dilutions were then prepared in 500 μL BBT+ buffer, and the sample was incubated overnight at 4°C. On the next day, the sample was washed with PBT+ buffer, then incubated with a pAG-MNase 1:100 dilution in 500 μL BBT+ for 4 hours at room temperature. For DNA cleavage, a Wash+C buffer was prepared by combining 1.5 mL Wash+ buffer with 30 μL 100 mM CaCl2, followed by resuspending the sample in 150 μL Wash+C buffer and incubating it for 45 minutes at 4°C. Then, a 2XSTOPyR buffer + was freshly prepared by adding 1600 μL 2XSTOP buffer and 10 μL RNaseA. At the end of the 45-minute incubation, 150 μL 2XSTOPyR buffer was added to the sample and incubated at 37°C for 30 minutes. The sample was centrifuged at 16,000 x g for 5 minutes, the supernatant was carefully extracted and transferred to a fresh Eppendorf tube. To this supernatant, 2 μL of 10% SDS and 2.5 μL of 20 mg/mL Proteinase K were added, and the mixture was thoroughly mixed using a brief vortexing procedure. Subsequently, the sample was incubated at 50°C in a water bath for a period of 2 hours. It’s important to note that this can be stopped at this step and the samples can be stored at −20 C. The magnetic beads and MXP were brought to room temperature before proceeding. 150 μL of the supernatant was extracted for subsequent bead cleanup, while keeping the remainder as a backup. 20 μL of AmpureXP bead slurry and 280 μL of MXP buffer were added to the sample and incubated for 15 minutes at room temperature. Using a magnetic rack, the beads were collected and incubated for 5 minutes. The supernatant was then discarded. The tubes were kept on the magnet and 1 mL of 80% ethanol was added to each tube without disturbing the beads. Using a magnetic rack, the beads were collected and incubated for 5 minutes. The supernatant was then discarded. The tubes were kept on the magnet and 1 mL of 80% ethanol was added to each tube without disturbing the beads. The sample was then incubated for a minimum of 30 seconds and the ethanol was gently aspirated to remove all traces of ethanol. While the tube remained on the magnet, the beads were air-dried for 2 minutes and resuspended in 10 μL of RNase and DNase free water. The samples were then incubated at room temperature for 2 minutes. Following this, the samples were kept on the magnet and the clear solution was transferred to a new Eppendorf tube. The DNA concentration was determined using a dsDNA high-sensitive Qubit assay and analyzed DNA size distribution in samples using a Fragment analyzer.

### DNA seq library preparation and analysis

The NEBNext Ultra II DNA Library Prep Kit for Illumina (E7645,E7103) protocol was followed for library preparation. Reads were first evaluated for their quality using FastQC (v0.11.8, RRID:SCR_014583).^99^ Reads were trimmed for adaptor sequences using Trim Galore! (v0.6.6, RRID:SCR_011847) and aligned to the dm6 reference genome version for drosophila melanogaster using Bowtie2 (version 2.2.8 RRID:SCR_016368) with parameters -q -I 50 -X 700 --very-sensitive-local --local --no-mixed --no-unal --no-discordant.^106^ Binary alignment maps (BAM) files were generated with samtools v1.9 ^107^ and were used in downstream analysis. MACS2 v2.1.0 ^108^ was used to call significant peaks for samples. IgG was used as control to call peaks. Peaks within ENCODE blacklisted regions and repetitive sequences larger than 100 bases were removed. Coverage tracks were generated from BAM files using deepTools 3.2.1^109^ bamCoverage function with parameters– normalize using RPKM–bin size 10. For genomic annotation promoters (−500 b to +500 b) relative to the TSS were defined according to the drosophila dm6 reference genome version. ChipSeeker (v1.36.0) was used to annotate Stonewall peaks. Stonewall binding motifs are called using HOMER (v4.10) findMotifsGenome function. Chromatin states are called using ChromHMM (v1.23). Heatmaps of genomic regions were generated with deepTools 3.2.1 computeMatrix and plotHeatmap commands, or EnrichedHeatmap (v1.30.0). PCA plot of histone modifications was generated using deepTools 3.2.1 multiBigwigSummary and plotPCA functions.

## Acknowledgments

We thank members of the Rangan and Dr. Miler T. Lee for their comments on the manuscript. We thank Julia Tang, Shruti Venkat and Paloma Bravo for helping with the HCR protocol. We thank Dr. Kahini Sarkar for helping with the cartoon. We thank the BDSC, VDRC, BDGP Gene Disruption Project, and Flybase for reagents and resources. P.R. is funded by the NIH/NIGMS (R01GM111779, RO1GM135628 and R56AG082906). This work was supported in part by the Bioinformatics for Next Generation Sequencing (BiNGS) shared resource facility within the Tisch Cancer Institute at the Icahn School of Medicine at Mount Sinai, which NIH grant P30CA196521 partially supports, the Black Family Stem Cell Institute, and the CDRB department at the Icahn School of Medicine at MSSM. This work was also supported in part through the computational resources and staff expertise provided by Scientific Computing at the Icahn School of Medicine at Mount Sinai and supported by the Clinical and Translational Science Awards (CTSA) grant UL1TR004419 from the National Center for Advancing Translational Sciences. The research reported in this paper was supported by the Office of Research Infrastructure of the National Institutes of Health under award number S10OD026880.

## Author contributions

Conceptualization, P.R. and N.M.K.; methodology, P.R. and N.M.K.; software and formal analysis, D.H. and G.K., and; investigation, D.H., G.K., and N.M.K; resources, P.R.; data curation, N.M.K., G.K., and.; writing – original draft, P.R. and N.M.K.; writing – review & editing, P.R,N.M.K, D.H., G.K. and M.J.; visualization, N.M.K.; supervision, P.R.; project administration, P.R.; funding acquisition, P.R.

---

**Supplementary Figure 1.**
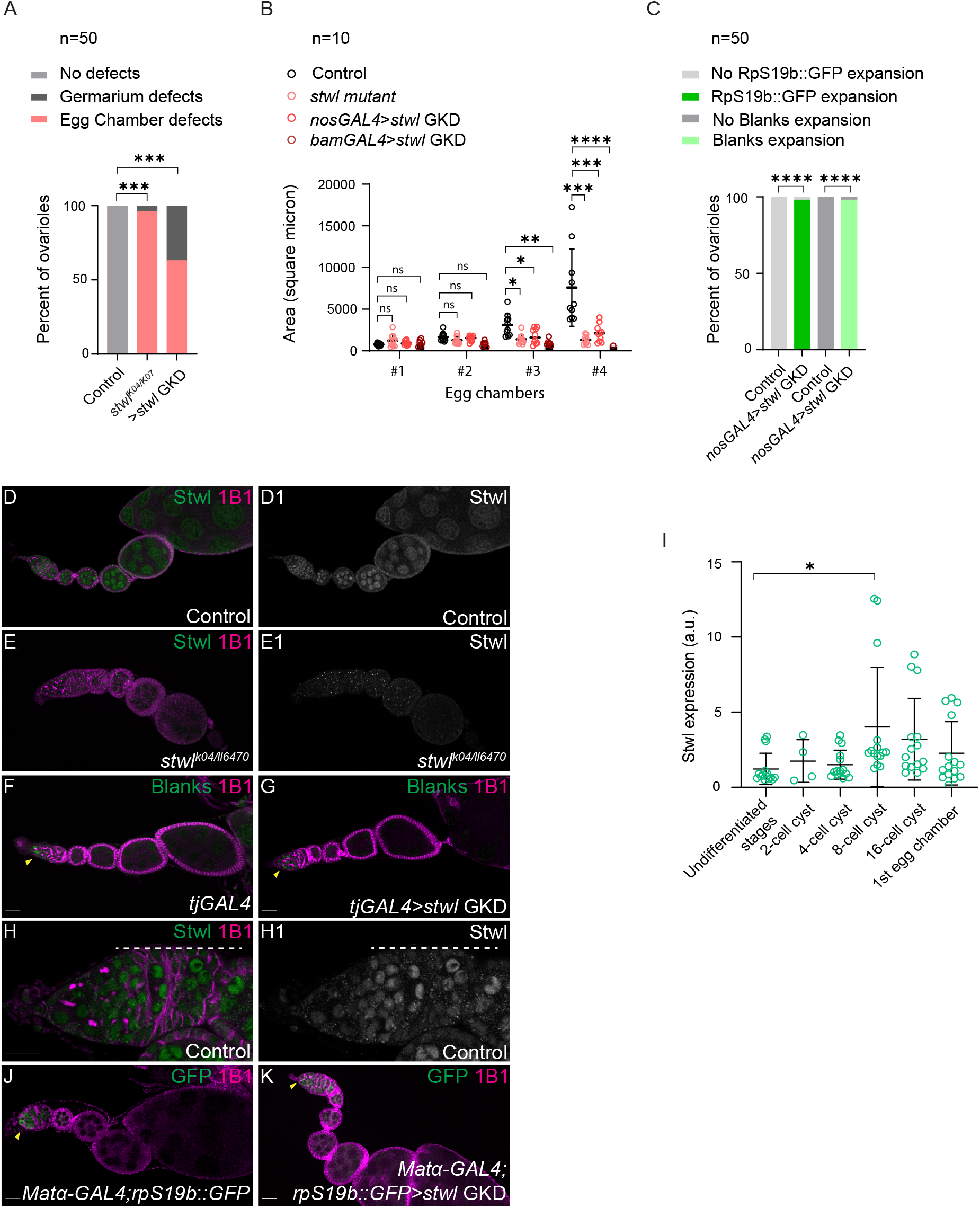
*stwl* is required for silencing the *rpS19b* reporter and Blanks during oogenesis. (A) Phenotypic defects quantification of *stwl* GKD and *stwl* mutants compared to control wild type (WT) ovaries. Statistics: Two tailed t-test; n = 50 ovarioles per genotype; ns, P> 0.05; ∗P < 0.05; ∗∗P < 0.01; ∗∗∗P < 0.001; ∗∗∗∗P< 0.0001. (B) Quantification of the size of the egg chambers of germline *stwl* GKD and *stwl* mutants compared to control WT ovaries. Graph shows that germline *stwl* GKD and *Stwl* mutants egg chambers do not grow in size compared to WT ovaries egg chambers that grow in size and lead to the formation of an egg. n=10. Statistics: Two tailed t-test; n = 50 ovarioles per genotype; ns, P > 0.05; ∗P < 0.05; ∗∗P < 0.01; ∗∗∗P < 0.001; ∗∗∗∗P < 0.0001. (C) Quantification of early oogenesis proteins RpS19b and Blanks expression in *stwl* GKD and *stwl* mutants compared to control WT ovaries. RpS19b::GFP and Blanks persist in egg chambers of *stwl* GKD and *stwl* mutants compared to the egg chambers of control. n=50. Statistics: Two tailed t-test; n = 50 ovarioles per genotype; ns, P > 0.05; ∗P < 0.05; ∗∗P < 0.01; ∗∗∗P < 0.001; ∗∗∗∗P < 0.0001. (D-D1) Ovariole of control *nosGAL4* ovaries (D) and in grayscale (D1) stained for Stwl (green, right in grayscale) and 1B1 (magenta). Stwl is expressed in the soma and germline of ovaries. (E-E1) Ovariole of *stwl^k^*^04^*^/ll^*^6470^ (E) and in grayscale (E1) stained for Stwl (green, right in grayscale) and 1B1 (magenta). Stwl expression is attenuated in *stwl^k^*^04^*^/ll^*^6470^ compared to the control. (F-G) Ovariole of control *TjGAL4* (F) and *TjGAL4>stwl* KD (G) stained for blanks (green) and 1B1 (magenta). *stwl* GKD in the soma did not result in any phenotypic defects and no Blanks expression in the differentiated egg chambers. (H-H1) Germarium of control *nosGAL4* ovaries (H) and in grayscale (H1) stained for Stwl (green,right in grayscale) and 1B1 (magenta). Stwl is expressed in soma and germline of ovaries and its levels increase during cysts stages. (I) Arbitrary unit (a.u.) quantification of Stwl expression in germaria of control ovaries (green). In the control, Stwl is expressed in the undifferentiated cells and its expression increases in the cysts stages. Statistics: Two-tailed t-test; n=5 ovarioles per genotype; ∗P < 0.05. (J-K) Ovariole of control *Matα-GAL4;rpS19b::GFP* (J) and *Matα-GAL4;rpS19b::GFP>stwl* GKD (K) stained for GFP (green) and 1B1 (magenta). RpS19b::GFP is expressed in undifferentiated stages in the control and upon *stwl* GKD in the differentiated stages using *Matα-GAL4*, no phenotypic defects are observed in the ovaries: egg chambers grow in size and GFP expression is attenuated in the egg chambers.

**Supplementary Figure 2.**
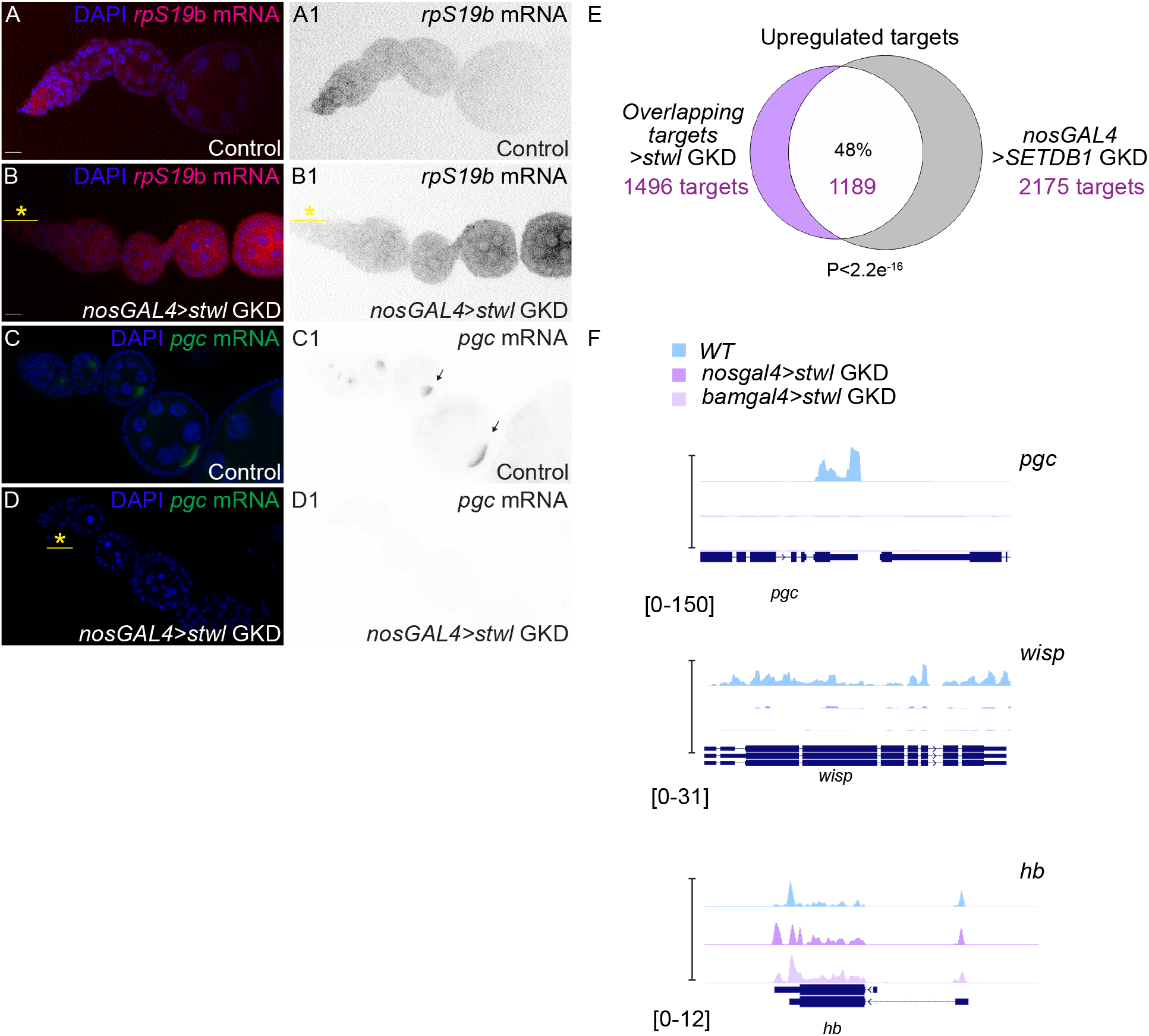
Stwl regulates the silencing of early oogenesis genes and activation of some maternal genes. (A–A1) Ovariole of control *nosGAL4* (A) and in grayscale (A1) probed for *rpS19b* RNA (red) and DAPI (blue). *rpS19b* RNA is expressed in the undifferentiated stages of the ovariole and is attenuated in the egg chambers. (B–B1) Ovariole of *nosGAL4> stwl* KD (B) and in grayscale (B1) probed for *rpS19b* RNA (red) and DAPI (blue). *rpS19b* RNA persists in the egg chambers of ovariole in contrast to the control where it is only expressed in the undifferentiated stages and is attenuated in the egg chambers.Yellow asterisk indicates loss of germline stem cells (GSCs) in *stwl* GKD. (C–C1) Ovariole of control *nosGAL4* (C) and in grayscale (C1) probed for *pgc* RNA (red) and DAPI (blue). *pgc* RNA localizes to the developing oocyte. (D–D1) Ovariole of *nosGAL4>stwl* KD (D) and in grayscale (D1) probed for *pgc RNA* (red) and DAPI (blue). *pgc* RNA is attenuated in the egg chambers of *stwl* GKD ovarioles in contrast to the control. (E) Venn diagram of overlapping upregulated genes from RNA-seq of *nosGAL4 & bamGAL4 > stwl* GKD ovaries and *nosGAL4> SETDB1* GKD compared with controls. 48% of targets are shared between both, suggesting that while Stwl and SETDB1 regulate a cohort of genes, Stwl regulates a subset of its targets through a different pathway than SETDB1.Statistics: hypergeometric test P<2.2e^-16^. (F) RNA-seq tracks showing that *pgc* and *wisp* are downregulated upon *stwl* GKD (*nosGAL4)* in dark purple and in the cyst stages (*bamGAL4)* in light purple. *hunchback (hb)* is not changing upon *stwl* GKD (*nosGAL4)* in dark purple and in the cyst stages (*bamGAL4)* in light purple.

**Supplementary Figure 3.**
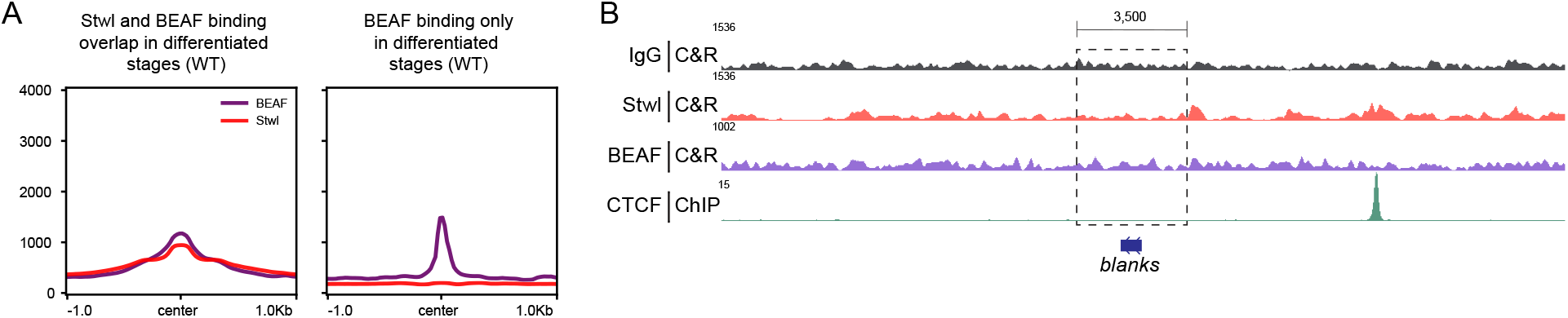
Stwl binds proximal to promoters and overlaps with boundary element proteins but does not directly regulate a large fraction of early oogenesis genes. (A) Profiles of CUT & RUN occupancy of Stwl peaks in differentiated stages of WT (red) ovaries overlapping with BEAF peaks occupancy in differentiated stages in WT ovaries (purple) and BEAF peaks occupancy in differentiated stages only in WT ovaries (purple). Profiles showing −1 Kb and +1Kb around the start and end of Stwl peaks shown in red. (B) CUT & RUN occupancy track for *blanks* locus for IgG (negative control) shown in grey, Stwl peaks in differentiated stages of WT ovaries (red), BEAF peaks in differentiated stages of WT ovaries (purple) and CTCF ChIP-seq peaks in larval CNSs (green) showing that Stwl, BEAF and CTCF are not flanking the upregulated target *blanks* from both sides as they are for other targets

**Supplementary Figure 4.**
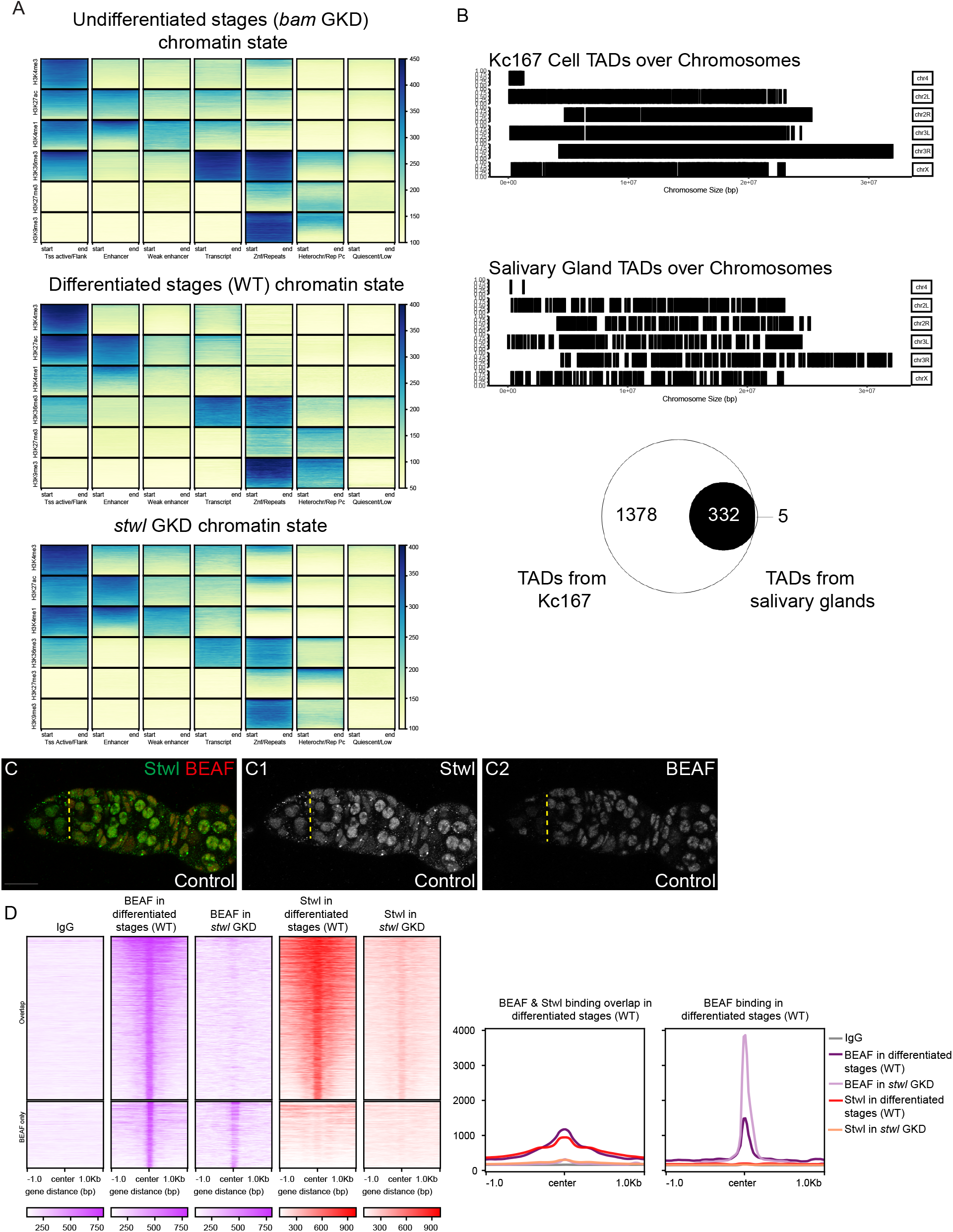
Stwl demarcates active and silenced genomic compartments. (A) Comprehensive chromatin state model of 7 states built from the combination of histone marks:H3K9me3, H3K27me3, H3K36me3, H3K4me1, H3K27ac and H3K4Me3 for undifferentiated stages (*bam* GKD), differentiated stages of oogenesis (WT ovaries) and in *stwl* GKD. (B) Maps of TADs on chromosomes of Kc167 cells and salivary glands. Venn diagram showing TADs of Kc167 cells encompassing TADs from salivary glands and shown to be conserved across different tissues. (C-C1-C2) Germarium of control (C) and in grayscale (C1-C2) stained for Stwl (green, right in grayscale C1) and BEAF (red, right in grayscale C2) showing BEAF expression is attenuated in the undifferentiated stages. (D) CUT & RUN profiles and occupancy of Stwl peaks in differentiated stages of WT (red) ovaries overlapping with BEAF peaks occupancy in differentiated stages of WT ovaries (purple) and BEAF peaks occupancy in *stwl* GKD. Heatmaps showing −1 Kb and +1Kb around the start and end of Stwl peaks shown in red. The occupancy heatmap is divided into two clusters: overlap between BEAF and Stwl in blue and BEAF alone in purple.

**Supplementary Figure 5.**
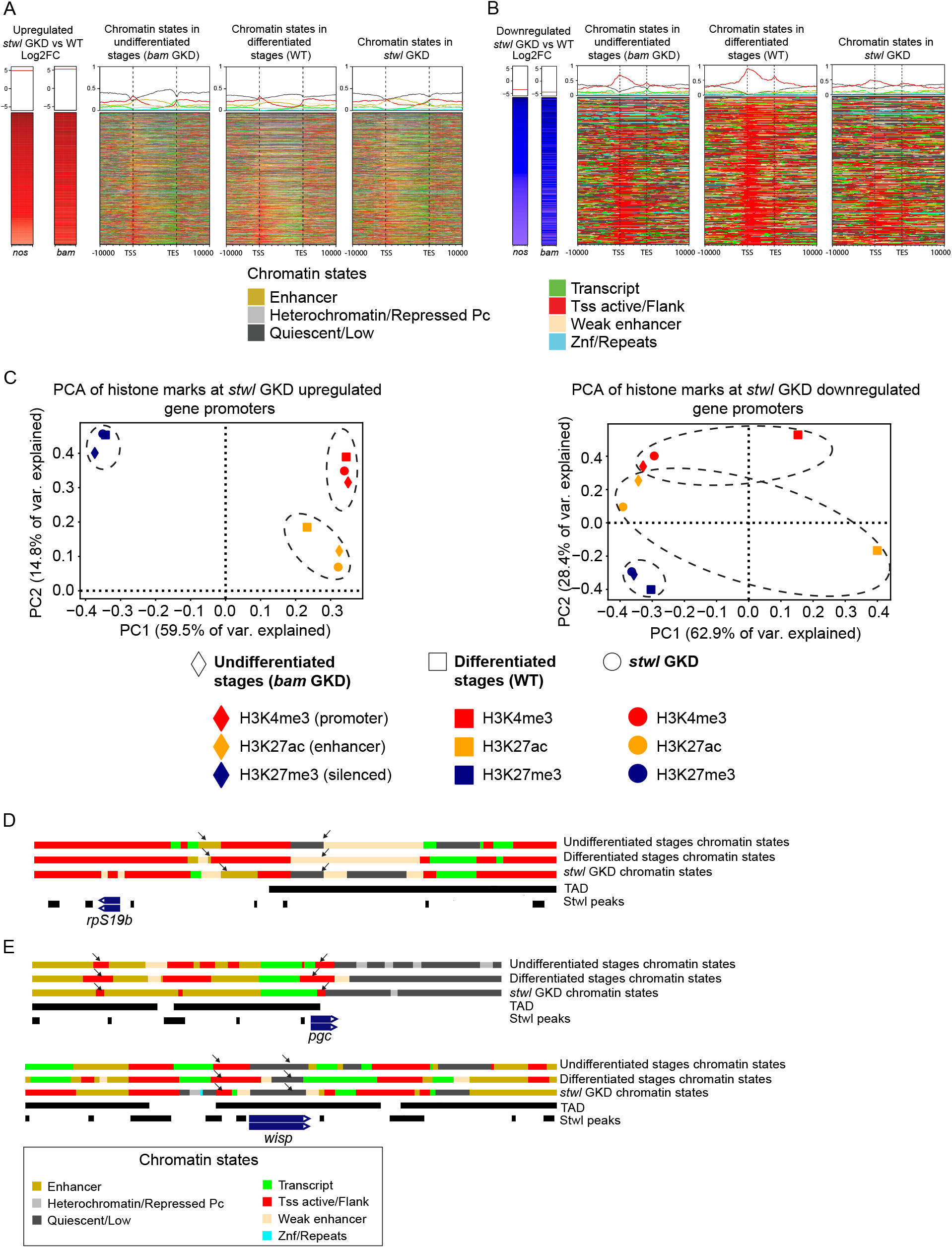
Stwl regulates enhancer activity to modulate gene expression. (A) Comprehensive analysis of 7 chromatin states of undifferentiated stages (*bam* GKD), differentiated stages (WT ovaries), and *stwl* GKD ovaries built using multiple histone marks shown for *stwl* GKD upregulated targets. Heatmap shown for targets as well as −10Kb and +10 Kb around the TSS and TES of genes. Different states marked by colors (gold: enhancer, light grey: heterochromatin/repressed, dark grey: quiescent/low, green: transcript, red: TSS active/flank, beige: weak enhancer and blue: Znf/repeats). This 7 state model shows enhancers changes at upregulated targets during differentiation which does not occur in *stwl* GKD. (B) Comprehensive analysis of 7 chromatin states of undifferentiated stages (*bam* GKD), differentiated stages (WT ovaries), and *stwl* GKD ovaries built using multiple histone marks shown for *stwl* GKD downregulated targets. Heatmap shown for targets as well as −10Kb and +10 Kb around the TSS and TES of genes. Different states marked by colors (gold: enhancer, light grey: heterochromatin/repressed, dark grey: quiescent/low, green: transcript, red: TSS active/flank, beige: weak enhancer and blue: Znf/repeats). This 7 state model shows active marks and quiescence changes at downregulated targets during differentiation which does not occur in *stwl* GKD. (C) PCA plots of histone marks H3K4me3 (red), H3K27ac (yellow) and H3K27me3 (navy) at *stwl* GKD upregulated and downregulated targets at gene promoters in undifferentiated stages (diamond shape), differentiated stages (square shape) and *stwl* GKD (circle shape). For upregulated targets H3K27ac in *stwl* GKD cluster close H3K27ac in the undifferentiated stages and further away from H3K27ac of the differentiated stages. For the downregulated targets, H3K4me3 and H3K27ac in *stwl* GKD cluster close to H3K4me3 and H3K27ac in the undifferentiated stages and further away from H3K4me3 and H3K27ac of the differentiated stages. (D) Chromatin states and TAD tracks of upregulated target *rpS19b* locus in undifferentiated stages, differentiated stages of WT ovaries and *stwl* GKD ovaries showing that *rpS19b* sits at TAD (black) boundary proximal to an active enhancer that decreases during differentiation. This decrease of the enhancer does not occur in *stwl* GKD (black arrows). Different states marked by colors (gold: enhancer, light grey: heterochromatin/repressed, dark grey: quiescent/low, green: transcript, red: TSS active/flank, beige: weak enhancer and blue: Znf/repeats). The genomic region proximal to *rpS19b* shows a decrease of active enhancers and an increase of weak enhancers during differentiation. These changes in enhancer states are attenuated in *stwl* GKD ovaries. (E) Chromatin states and TAD tracks of downregulated target *pgc* and *wisp* loci in undifferentiated stages, differentiated stages of WT ovaries and *stwl* GKD ovaries showing that *pgc* sits at TAD (black) boundary and that upon loss of *stwl*, weak enhancer (beige) flanking *pgc* is lost (black arrows). Different states marked by colors (gold: enhancer, light grey: heterochromatin/repressed, dark grey: quiescent/low, green: transcript, red: Tss active/flank, beige: weak enhancer and blue: Znf/repeats) *pgc and wispy*, downregulated genes, show an increase in active marks during differentiation and a decrease in quiescence. The decrease in quiescence and increase in the active marks is attenuated upon *stwl* GKD.

**Supplementary Figure 6.**
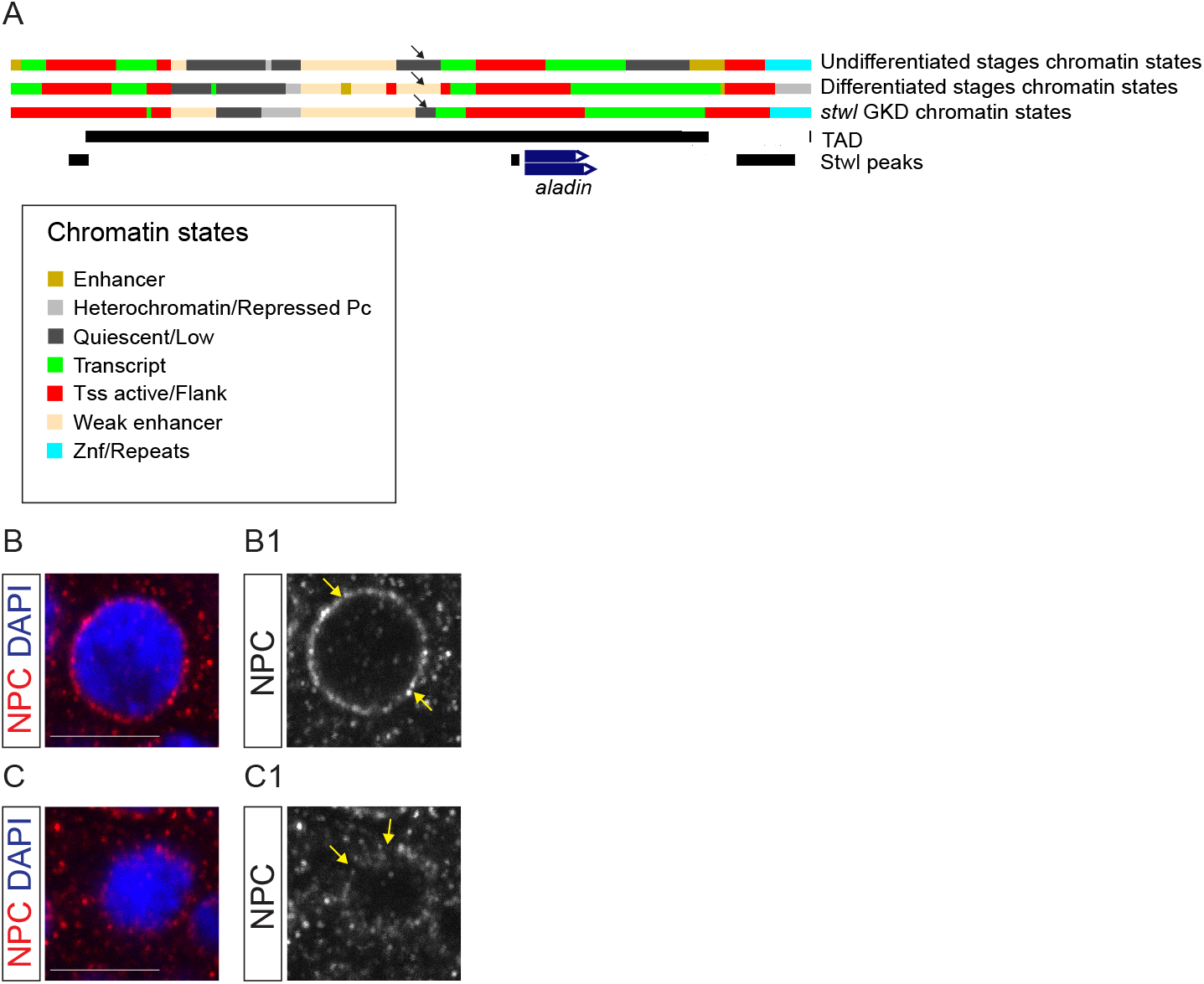
Stwl promotes the expression of nucleoporins to promote NPC formation. (A) Chromatin states and TAD tracks of the Nup *aladin* locus in undifferentiated stages, differentiated stages of WT ovaries and *stwl* GKD ovaries showing that *aladin* sits on TAD boundary (black), (black arrows). Nup *aladin,* which is downregulated upon the loss of *stwl*, in the genomic region proximal to it, there is a decrease of quiescence, and gain of a weak enhancer and active enhancer during differentiation. This change in quiescence and enhancer state does not occur upon *stwl* GKD. Different states marked by colors (gold: enhancer, light grey: heterochromatin/repressed, dark grey: quiescent/low, green: transcript, red: TSS active/flank, beige: weak enhancer and blue: Znf/repeats). (B-B1) Nurse cell nucleus of Control (B) and in grayscale (B1) stained for NPC in red (right grayscale B1) and DAPI in blue showing ring structure of NPC. (C-C1) Nurse cell nucleus of *stwl* GKD (C) and in grayscale (C1) stained for NPC in red (right grayscale C1) and DAPI in blue showing gaps in ring structure of NPC. Scale bars: 15 μm

**Supplementary Figure 7.**
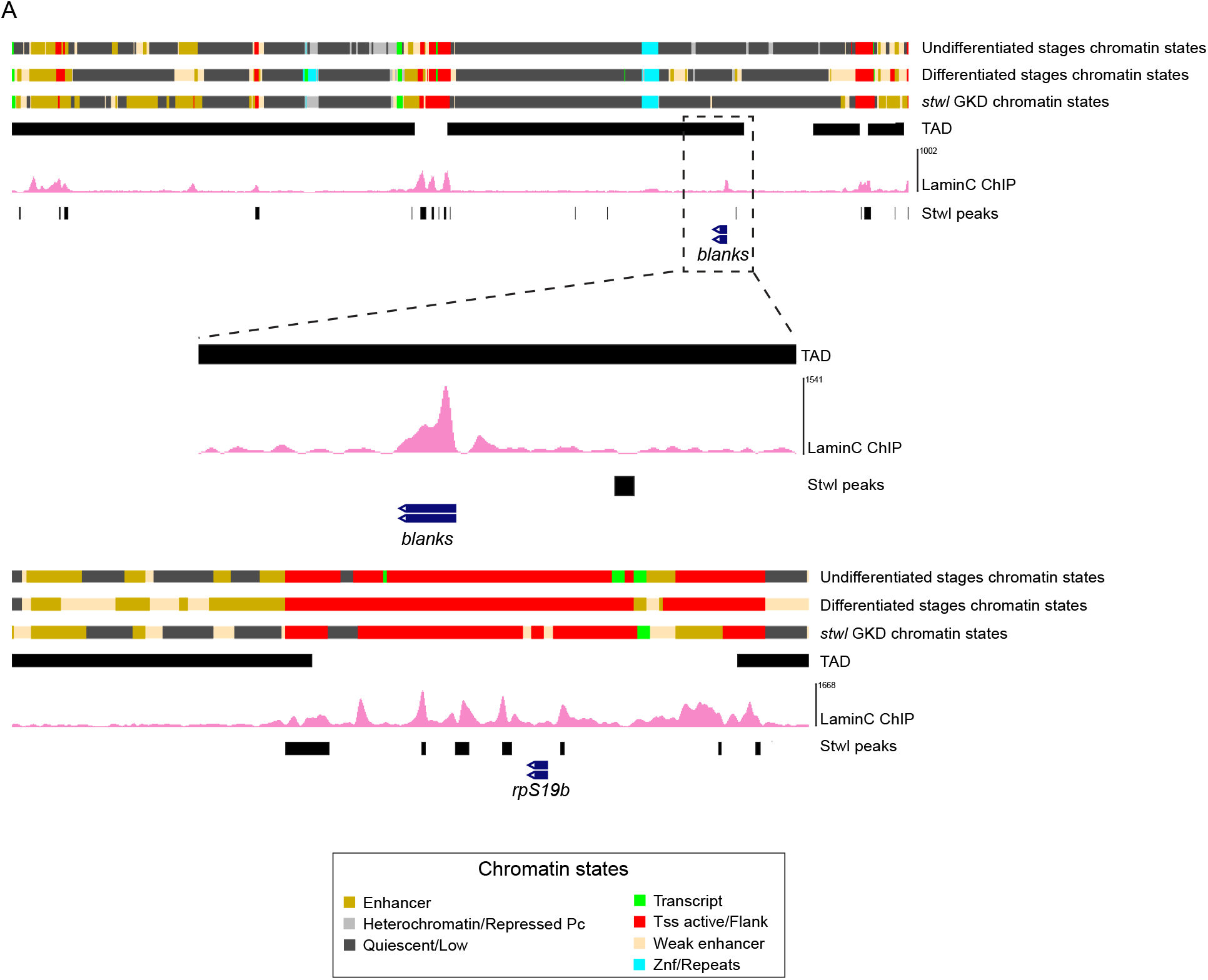
Early oogenesis genes are tethered to the nuclear periphery. (A) CUT & RUN occupancy tracks at the upregulated targets *rpS19b* & *blanks* loci for Stwl peaks in differentiated stages of WT ovaries (black) and Lamin C peaks from S2 cells ChIP (pink). Stwl peaks and Lamin C are observed on *rpS19b* locus from both sides and on both sides of TAD containing *blanks.* Different states are marked by colors (gold: enhancer, light grey: heterochromatin/repressed, dark grey: quiescent/low, green: transcript, red: Tss active/flank, beige: weak enhancer and blue: Znf/repeats).

